# The ATR kinase of *Trypanosoma brucei* links DNA damage signalling and monoallelic control of surface antigen gene expression during antigenic variation

**DOI:** 10.1101/435198

**Authors:** Jennifer Ann Black, Kathryn Crouch, Leandro Lemgruber, Craig Lapsley, Nicholas Dickens, Jeremy C. Mottram, Richard McCulloch

## Abstract

To evade mammalian immunity, *Trypanosoma brucei* switches the variant surface glycoprotein (VSG) expressed on its surface. Key to this reaction are controls exerted to ensure only one of many subtelomeric multigene VSG expression sites are transcribed at a time. DNA repair activities have to date been implicated only in catalysis of VSG switching by recombination, not transcriptional control. However, how VSG switching is signalled to guide the appropriate reaction, or to integrate switching into parasite growth, is unknown. Here we show that loss of ATR, a DNA damage signalling protein kinase, is lethal and causes increased nuclear genome lesions. ATR depletion also causes expression of mixed VSGs on the cell surface, increased transcription of genes from silent expression sites, and altered localisation of RNA Polymerase I and VEX1, factors involved in VSG transcription. The work therefore reveals that VSG expression control is mediated by a nuclear DNA damage signalling factor.

## Introduction

Multiple pathways and activities have evolved to tackle the wide range of stresses faced by cells. Damage to the genome is a one such stress and dealing with the myriad lesions that affect the genome is needed to ensure faithful transmission. A key, early step in execution of the eukaryotic DNA damage response is recognition and signalling of DNA lesions, a reaction in which phosphatidylinositol 3-kinase-related kinases (PIKKs) play a central role. Three DNA damage sensing PIKKs have been described: the DNA-dependent protein kinase catalytic subunit (DNA-PKcs), ataxia-telangiectasia mutated (ATM), and ataxia telangiectasia and Rad3-related (ATR) kinases. Each PIKK is recruited to damaged DNA by distinct binding partners, which provide recruitment to specific lesions and leads to activation of specific repair pathways (Lovejoy and Cortez, 2009). Indeed, interaction with a regulatory co-factor is a common mode of action for wider PIKKs, including SMG-1 (suppressor of morphogenesis in genitalia) and TOR (target of rapamycin)(Baretic and Williams, 2014). DNA-PKcs and ATM are both recruited to DNA double strand breaks (DSBs), but while the former protein kinase is targeted to this lesion by the Ku heterodimer and directs repair by non-homologous end-joining (NHEJ), the latter is recruited by the Mre11-Rad50-Xrs2 (MRX) complex and directs homologous recombination (HR) repair. ATR, in contrast, is recruited to single-stranded DNA coated by replication protein A (RPA) through binding to ATRIP (ATR interacting protein)(Wang et al., 2017) and, in mammals, by ETAA1 (Zou, 2017). Activation of ATR-ATRIP requires further recruitment of TopBP1 and the Rad9-Hus1-Rad1 complex. Single-stranded DNA can form in many settings, meaning ATR has been implicated in repair of DSBs and intra-strand crosslinks (Sirbu and Cortez, 2013), and in telomere homeostasis (Maciejowski and de Lange, 2017). However, damage signalling by ATR is most intimately linked with replication stress, where the PIKK stabilises replication forks that encounter impediments to their passage, such as damage, DNA secondary structures, the transcription machinery and RNA-DNA hybrids (Saldivar et al., 2017; Zeman and Cimprich, 2014). ATR’s role in the replication stress response is to limit replication fork collapse, thereby allowing the stalled replisome to resume DNA synthesis, which involves regulation of cell cycle progression, co-ordinating usage of sites of DNA replication initiation (termed origins) and modulation of replisome activity.

*Trypanosoma brucei* is one of several causative agents of African trypanosomiasis, which can affect both humans and domesticated animals (Morrison et al., 2016). All salivarian trypanosomes are extracellular parasites and survive elimination by the adaptive immune response of mammals though stochastic changes in their protective Variant Surface Glycoprotein (VSG) coat. Such surface antigen switching, termed antigenic variation, is widespread in bacterial, viral and eukaryotic pathogens but appears to have evolved remarkable mechanistic complexity in *T. brucei*. In any given cell only a single *VSG* gene is normally actively transcribed to generate a homogenous VSG coat (Manna et al., 2014). VSG transcription in the mammal occurs in telomeric bloodstream VSG expression sites (BES), of which around 15 are present (Berriman et al., 2002; Hertz-Fowler et al., 2008). Mechanisms have therefore evolved to ensure that only one BES is actively transcribed, and that such monoallelic expression is flexible enough to allow transcription of the active BES to be silenced with concomitant activation of previously silent BES, thereby eliciting a VSG coat switch. The active ES is transcribed by RNA Polymerase (Pol) I and localises to an extra-nucleolar body (the expression site body, ESB) in the *T. brucei* nucleus (Lopez-Farfan et al., 2014; Navarro and Gull, 2001). Perturbation of a number of processes have been revealed to undermine monoallelic expression, including telomere (Jehi et al., 2014a; Jehi et al., 2016; Yang et al., 2009) and nuclear envelope integrity (DuBois et al., 2012; Maishman et al., 2016), chromatin status (Alsford and Horn, 2012; Aresta-Branco et al., 2016; Denninger et al., 2010; Hughes et al., 2007; Narayanan and Rudenko, 2013; Povelones et al., 2012), chromatid cohesion (Landeira et al., 2009) and inositol phosphate signalling (Cestari and Stuart, 2015). In addition, potentially kinetoplastid-specific monoallelic control factors are present, such as VEX1 (Glover et al., 2016). Understanding how VSG transcriptional switching, which appears to be a co-ordinated process (Chaves et al., 1999), is executed (Figueiredo et al., 2008), initiated (Batram et al., 2014) and signalled (see below) has been less studied.

A further route for VSG switching is recombination of a silent VSG into the BES (McCulloch et al., 2015), using a genomic archive numbering >1000 VSG genes and pseudogenes (Berriman et al., 2005; Cross et al., 2014). Extensive evidence indicates that HR, catalysed by RAD51 (McCulloch and Barry, 1999) and mediated by further factors (Devlin et al., 2016b; Dobson et al., 2011; Hartley and McCulloch, 2008; Kim and Cross, 2010, 2011; Proudfoot and McCulloch, 2005; Trenaman et al., 2013), directs the switching of functionally intact *VSG*s. It is less clear how VSG pseudogenes are recombined, but the flexible, combinatorial assortment of these sequences generates huge levels of expressed VSG diversity, particularly in chronic infections (Hall et al., 2013; Marcello and Barry, 2007; McCulloch and Field, 2015; Mugnier et al., 2015). A final mechanistic complexity is found in emerging evidence that VSG switching may be intricately linked to density-dependent signalling that controls cell and life cycle progression of trypanosomes (Batram et al., 2014; Macgregor et al., 2011; MacGregor et al., 2012; Matthews et al., 2015). As for transcriptional switching, the trigger for VSG switching by recombination is still being sought, with BES DSBs (Boothroyd et al., 2009; Glover et al., 2013a), BES replication (Benmerzouga et al., 2013; Devlin et al., 2016a; Devlin et al., 2016b) and telomere shortening (Hovel-Miner et al., 2012) having been suggested.

The signalling events that respond to triggers of VSG switching have not been characterised, though loss of MRE11 (from the MRX complex) appears to have little impact (Robinson et al., 2002). Understanding how VSG switching is signalled would take us closer to revealing the nature of such a trigger, including the form of DNA lesion that might direct VSG recombination, whether switching is linked to genome replication, and how the reaction might be linked more broadly to the cell cycle. To date no work has examined whether PIKKs contribute to these decision points or, indeed, to antigenic variation at all. Here we show that loss of ATR in mammal-infective *T. brucei* cells results in rapid growth impairment and heightened sensitivity to a wide range of DNA damage, consistent with an essential role in genome maintenance. In addition, loss of ATR rapidly undermines ES expression control, seen as reduced expression of the singularly expressed VSG RNA and protein, coupled with increased transcription from potentially all silent BES and expression of mixed VSG coats. These BES transcription changes after ATR RNAi not only closely resemble the effects seen after VEX1 RNAi, but ATR loss leads to altered localisation of VEX1 and RNA Pol I. Thus, we reveal a mechanistic link between DNA damage signalling and monoallelic control of VSG expression during *T. brucei* immune evasion.

## Results

### ATR is essential for *T. brucei* proliferation and for survival following DNA damage

A putative homologue of the ATR kinase has previously been identified in *T. brucei* (Parsons et al., 2005) and preliminary RNAi analysis revealed impaired proliferation of bloodstream form (BSF) *T. brucei* cells (Jones et al., 2014). However, several proteins involved in mediation of ATR activity have yet to be identified in *T. brucei*, including ATRIP and the downstream target CHK1 (checkpoint kinase 1)(Goto et al., 2015). A homologue of TopBP1 has been predicted (Genois et al., 2014) but not validated. The 9-1-1 complex plays important, novel roles in *Leishmania* genome maintenance (Damasceno et al., 2013; Damasceno et al., 2016; Nunes et al., 2011) but interaction with ATR directly or indirectly has not been assessed, and no work has examined 9-1-1 function in *T. brucei*. Thus, how (and if) *T. brucei* ATR (TbATR) acts in damage signalling, including conservation of its associated machinery, is unknown.

To examine the effect of TbATR loss, *in vitro* proliferation of BSF cells after tetracycline (Tet) induced RNAi was examined in two clones, one expressing the kinase from its own locus translationally fused to 12 copies of the myc epitope at the C-terminus (TbATR-12myc). In both clones growth ceased from 24 hrs (Fig.1A).

**Figure 1.**
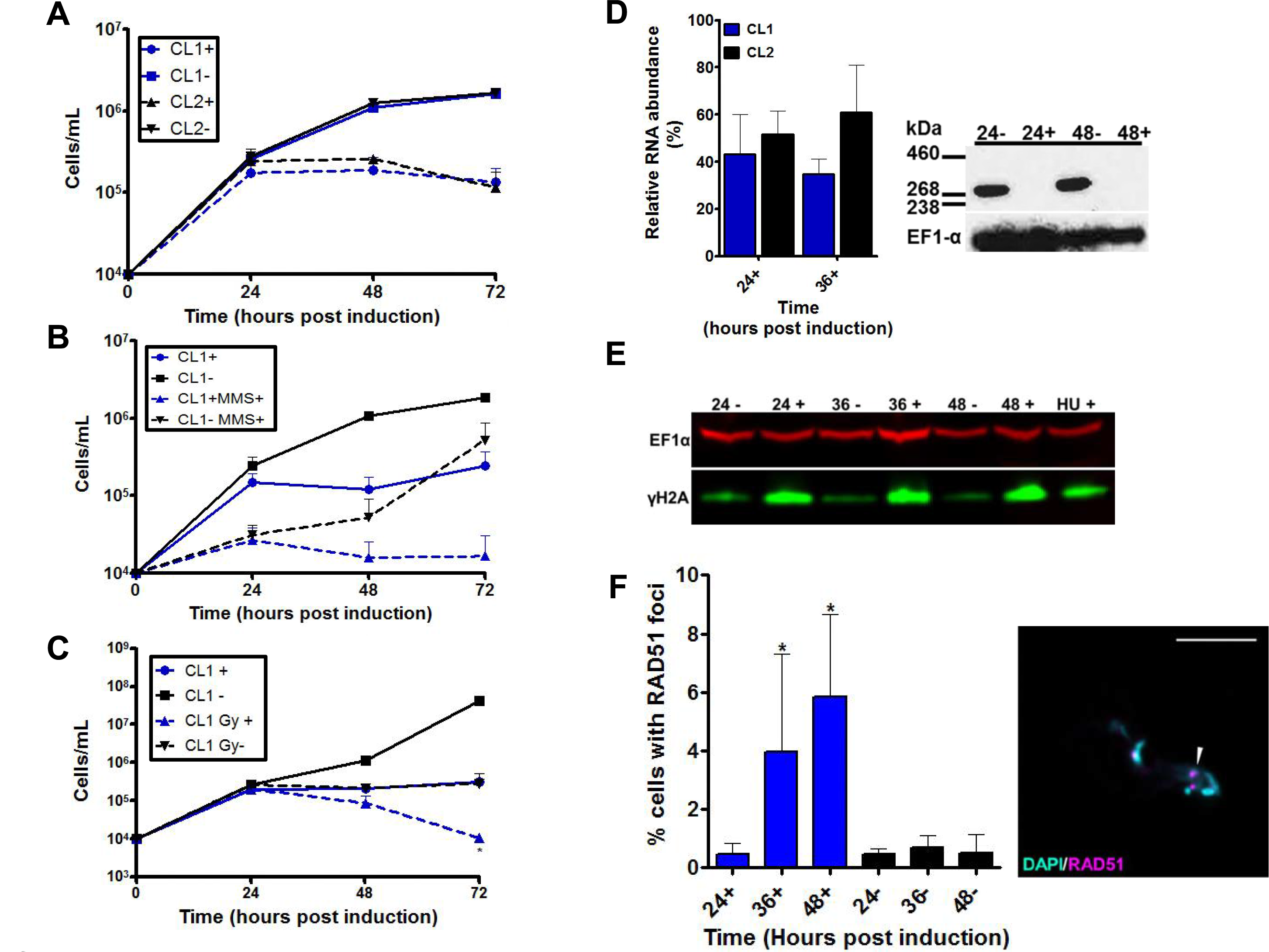
Loss of ATR halts growth of bloodstream form *T. brucei* and impairs the parasite’s ability to tackle nuclear genome damage. (A) The upper graph shows cell density following TbATR RNAi after addition of tetracycline (+, dashed lines) over 72 hours, or when RNAi is not induced (-, solid lines); error bars show ± SEM from three experiments, and two clones (CL1, blue, and CL2, black) are compared. Growth curves showing cell density over time with and without TbATR RNAi induction in the absence or presence of (B) Methylmethanosulfonate (MMS, 0.0003%) and (C) ionising radiation (IR, 150 Gy). Data are shown for CL1 only; error bars denote ± SEM (n=3). (D) The graph shows TbATR RNA abundance assessed by RT-qPCR in two clones (CL1, CL2) 24 and 36 hrs after RNAi relative to uninduced cells (set at 100%) at the same times; error bars show ± SEM (n=3 for CL1, and n=2 for CL2). Abundance of TbATR-12myc in CL2 is shown by western blot after 24 or 48 hrs growth with and without addition of Tet (+ and −, respectively); TbATR-12myc (∼270 kDa) was detected using anti-myc antiserum, and anti-EF1α acts as a loading control. (E) Expression levels of the phosphorylated variant of histone H2A (γH2A, green) are shown by western blotting of whole cell extracts after 24, 36 or 48 growth with (+) and without (−) addition of Tet; EF1α (red) serves as a loading control, and levels are shown after 48 hrs growth of uninduced cells in the presence (+) of 0.06 mM HU (blots are shown only for CL2). (F) Quantification of the percentage of cells in the population that harbour RAD51 foci after 24, 36 or 48 hrs growth with (+) and without (−) RNAi induction. Error bars show ± SEM (n = 3, >200 cells counted/experiment); (*) = p<0.05, Mann Whitney U test. Representative image of a tet+ cell harbouring RAD51 foci (white arrow) is show to the right of the graph. N- and k-DNA is DAPI stained (cyan) and anti-RAD51 antiserum was used to detect RAD51 foci (magenta). Scale bar ; 5 µm.

RT-qPCR of both clones and western analysis of the TbATR-12myc expressing clone (Fig.1D) showed growth impairment was accompanied by reduced levels of TbATR RNA and loss of detectable myc-tagged protein from 24 hrs after RNAi induction. To ask if TbATR plays a role in the DNA damage response, we examined whether its loss sensitises BSF cells to genotoxic stress by evaluating growth in the presence of methyl methanesuplhonate (MMS, an alkylator; Fig.1B) or hydroxyurea (HU, a ribonucleotide reductase inhibitor; Fig.1 – Figure supplement 1A), or after exposure to ionising radiation (Fig.1C) or UV (a nucleic acid cross-linker; Fig.1 – Figure supplement 1A). Uninduced cells grew more slowly at these concentrations of MMS or HU, and after exposure to ionising radiation. In all cases, and in both RNAi clones (Fig.1 – Figure supplement 1B), growth retardation was more marked after RNAi and damage. Exposure of uninduced cells to 1500 J/m^2^ UV had little effect on proliferation, but growth was markedly slowed after TbATR RNAi relative to the RNAi induced untreated cells (Fig.1 – Figure supplement 1A,B). Taken together, these data indicate that loss of TbATR sensitises BSF cells to a range of genotoxic agents, suggesting the PIKK contributes to the response of *T. brucei* to a variety of DNA lesions.

### Loss of ATR leads to nuclear DNA damage

To ask if the above phenotypes reflect nuclear roles for TbATR, we tested whether or not loss of the PIKK causes discernible genome damage. Phosphorylation of histone H2A on Thr130 has been described in *T. brucei* (Devlin et al., 2016b; Glover and Horn, 2012) and in *L. major* (Damasceno et al., 2016) after exposure to different genotoxic stresses or repair gene mutation, and thus represents a kinetoplastid variant of the γH2A(X) damage-response nuclear chromatin modification (Biterge and Schneider, 2014). In uninduced TbATR RNAi cells anti-γH2A antiserum recognised some protein in western blots (Fig.1E), and IF signal could be seen in the nucleus of some cells (Fig.1 – Figure supplement 2A). Nonetheless, western blotting showed that levels of γH2A increased 24 hrs after TbATR RNAi (Fig.1E, Fig.1 – Figure supplement 2B), while IF showed that both the number of cells with γH2A nuclear signal and the signal intensity increased after RNAi (Fig.1 – Figure supplement 2A). Thus, loss of the PIKK results in increased nuclear genome damage. To probe the nuclear damage further, we examined localisation of RAD51, an HR enzyme that forms nucleoprotein filaments on single stranded DNA, including at DSBs, and relocalises to discrete nuclear foci in *T. brucei* after phleomycin treatment or induction of a DSB by ISceI (Devlin et al., 2016b; Dobson et al., 2011; Glover and Horn, 2014; Trenaman et al., 2013). IF with anti-RAD51 antiserum showed that, in keeping with previous work, only a small (<1%) part of the uninduced cell population had detectable nuclear foci (Fig.1F, Fig.1 – Figure supplement 2C). However, 36 and 48 hrs after TbATR RNAi a significant increase in the number of cells with RAD51 foci was seen (Fig.1F, ∼6% of the population), indicating at least some of the nuclear damage resulting from loss of TbATR is recognised by RAD51.

### Altered VSG expression site transcription emerges rapidly after ATR RNAi

To ask if TbATR loss results in altered gene expression total RNA was prepared, in triplicate, after 24 hrs of RNAi induction and subjected to RNAseq, comparing changes in gene-specific read abundance relative to uninduced cells (Fig.3). To map sequence reads not only to the core genome but also to specific VSG BES (which share sequence homology amongst the ESAGs) MapQ filtering was applied (Hutchinson et al., 2016). 231 transcripts (including TbATR) were significantly differentially expressed in the RNAi induced cells relative to uninduced (Fig.2A, Table S1). 80% of these genes showed increased RNA levels, with VSG and ESAG genes from the silent BES being the most abundant category (∼35% of the total; Fig.2B) and showing the greatest increases (Table S1; 1.5-23 fold). Also prominently represented (∼19%; 1.3-4.8 fold increase) were other putative RNA Pol I-transcribed genes, including VSG or VSG-related genes from outwith the BES, and procyclin or procyclin-associated genes (PAGs) that are normally expressed in tsetse-infective *T. brucei* cells. Notably, only three RNA Pol I-transcribed genes displayed reduced expression (Fig.2B): two ESAGs from the active BES (BES1; Table S1; -1.3 fold) and one PAG (Table S1; -1.3 fold). Taken together, these data indicate that the earliest and most pronounced effect of TbATR loss is altered expression of RNA Pol I genes, which appears to mainly result from increased transcription from the silent BES. GO term enrichment analysis confirmed this interpretation: the most pronounced enrichment (in terms of number of genes affected and level of enrichment) was in the up-regulated cohort (Table S2) and in functions associated with the VSG and BES (Fig.2C), including evasion of the host immune response and antigenic variation (biological function), and cell surface or membrane (cellular location).

**Figure 2.**
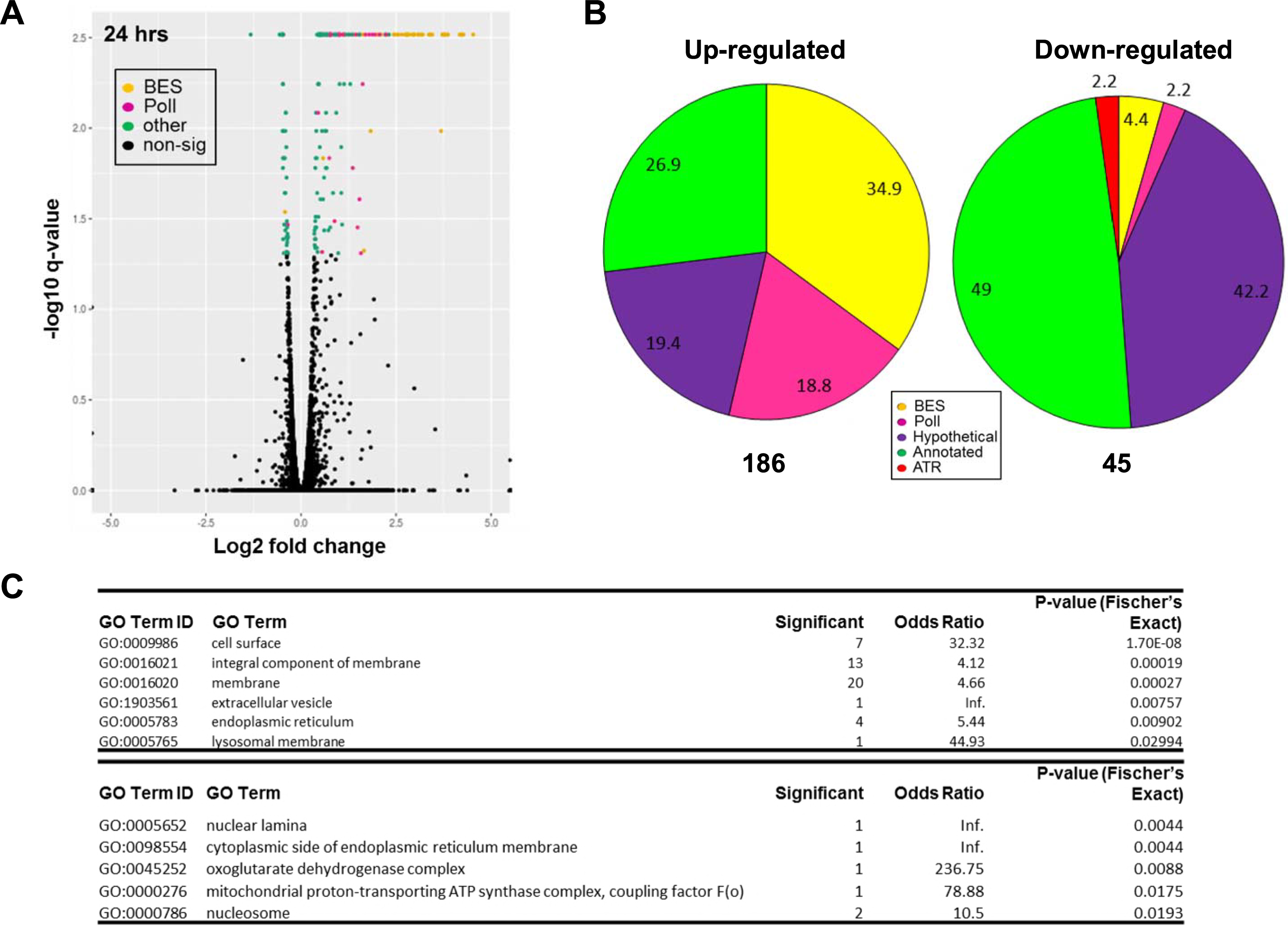
ATR RNAi leads to de-repression of surface antigen gene expression in bloodstream form *T. brucei* cells. (A) Volcano plots showing transcripts differentially expressed 24 hours after TbATR RNAi in BSF cells relative to uninduced controls. Log10 transformed p-values for each gene identified by RNAseq are plotted against the log2 transformed fold-change in FPKM (fragments per kilobase transcript per million bases, determined by CuffLinks). Data are averages from three biological replicates and transcripts are annotated as follows: BES (yellow), from a bloodstream expression site; Pol I (pink), transcribed by RNA Pol I but not in BES; other (green), non-Pol I-transcribed gene; non-sig (black), non-significant change in abundance. (B) Pie charts summarising differentially expressed genes (left, decreased transcript abundance; right, increased transcript abundance) 24 hours after RNAi induction relative to uninduced cells. The number of genes in each category is expressed as a percentage of the total gene number (shown below each chart). BES genes and RNA Pol I transcribed genes are coloured yellow and pink, respectively; non-RNA Pol I genes with functional annotations (available on TriTrypDB) are shown in green, while those annotated as hypothetical are in purple; TbATR is shown in red. (C) Summary of GO terms significantly enriched in the differentially expressed gene cohort relative to the expected number of genes in the genome; enriched GO IDs and GO terms are shown for up-regulated genes under the headings Biological Function and Cell Location; gene IDs, down-regulated genes and enriched terms in heading Molecular Function are provided in Table S2.

**Figure 3.**
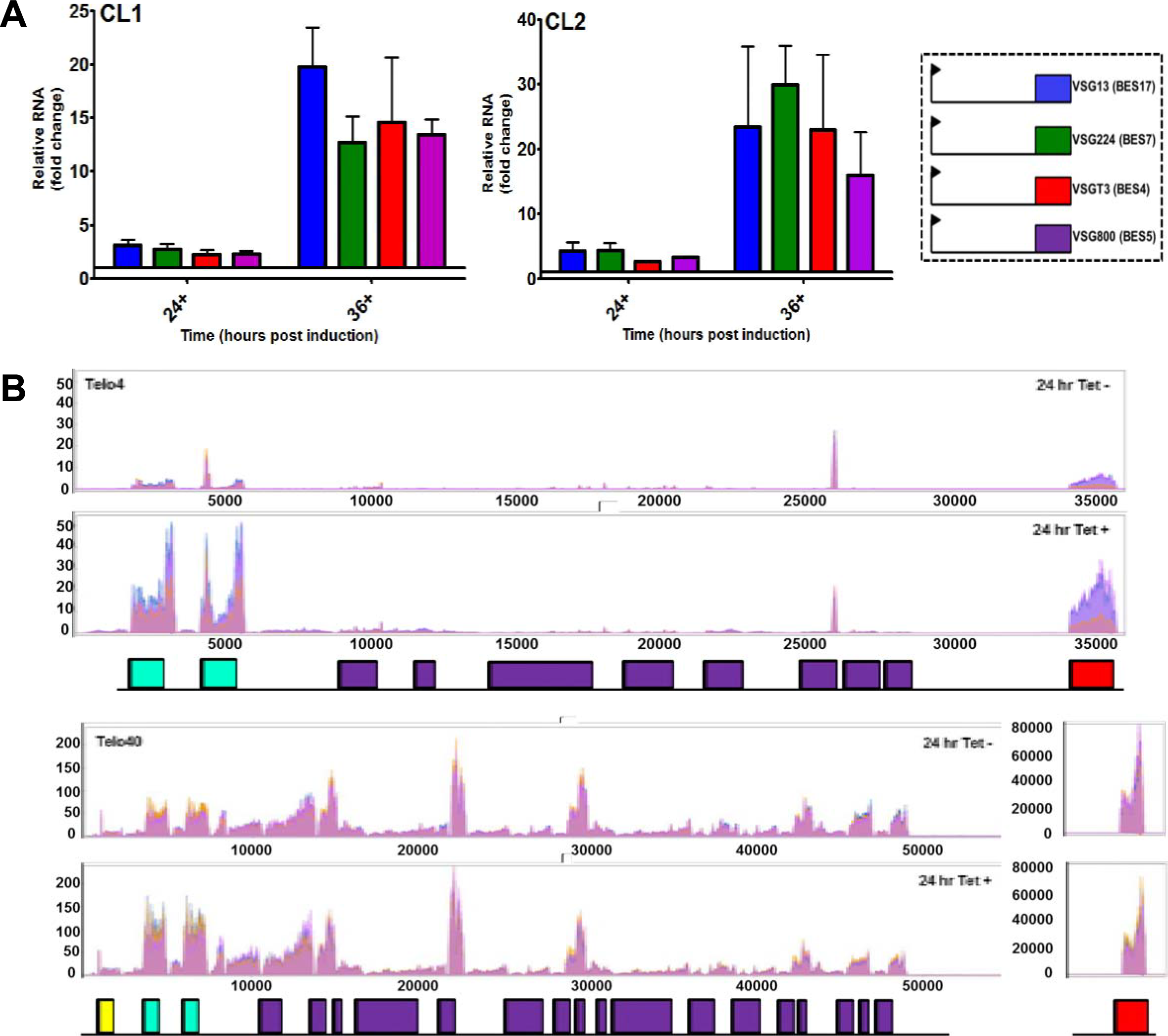
Loss of ATR impairs control of *T. brucei* bloodstream form VSG expression site transcription. (A) RNA levels of VSG genes, assessed by RT-qPCR, within four silent BES are shown (24 and 36 hours post RNAi) as fold-change relative to RNA levels uninduced cells. Data are shown for two clones (CL1, CL2) and error bars denote ± SEM (n=3 for CL1, n = 2 for CL2); VSGs are named as detailed in (Hertz-Fowler et al., 2008). (B) Mapping of MapQ filtered RNAseq reads is shown across a silent VSG bloodstream VSG expression sites (Telo4) and the single active site (Telo40) after 24 hrs growth with (T+) or without (T−) induction of TbATR RNAi; y axes show the number of reads that map relative to location in the transcription units, and data from three replicates are overlaid. ESAG6 and ESAG7 genes are shown in green, all other ESAGs are in purple, the VSG is in red, and a gene encoding resistance to hygromycin is in yellow. For Telo40, read mapping to the VSG is shown on a different scale to the rest of the expression site.

Of the 128 genes that displayed significantly differentially expressed transcripts and were not associated with RNA Pol I transcription (Fig.2B, Table S1), less bias towards increased levels and smaller changes were seen (86 increased 1.3-5 fold; 42 decreased 1.3-2.5 fold). Amongst those with functional annotations, GO term enrichment analysis (Table S2) did not reveal any activities represented by more than 5 genes (indeed most were only represented by a single gene). Thus, at this early stage after RNAi, RNAseq does not reveal potential pathways of core gene expression changes that might reveal functions regulated by TbATR to enact its putative signalling functions.

### ATR RNAi leads to loss of monoallelic VSG expression

Given that RNAseq suggests TbATR RNAi has the most pronounced impact on expression of the BES, we tested this prediction in more detail. Only a single VSG is normally expressed in a BSF cell and, in the strain used here, monoallelic expression results in the predominant expression of VSG221 (VSG2) from BES1 (Telo40; one of 14 characterised BES that contain distinct telomere-proximal VSGs)(Hertz-Fowler et al., 2008). RNAseq analysis suggested TbATR loss results in altered expression of most silent VSGs, since significantly increased RNA levels were detected for nine normally silent VSGs (1.7-7.9 fold; Table S1) 24 hrs after RNAi induction. To check the RNAseq, we performed RT-qPCR of four silent BES VSGs and confirmed increased levels of each in TbATR RNAi induced cells relative to controls (Fig.3A), with reasonable correspondence in fold-changes (2.2-3.1). RT-qPCR of the same VSGs 36 hrs after RNAi induction indicated even greater increases in abundance (12-30 fold; Fig.3A). Given this finding, and the RNAseq description of significantly reduced levels of two ESAG transcripts (1 and 3) from BES1 (telo40), allied to significant increases in ESAG transcripts from several silent BES at 24 hrs (Table S1), we performed RNAseq (again in triplicate, using the two RNAi clones) after 36 hrs RNAi and mapped the data from both 24 and 36 hrs using MapQ filtering to all BES (Fig.3B, Fig.3 – supplementary figures 1 and 2). The mapping revealed two main things.

First, read mapping indicated that levels of VSG221 (VSG2) transcript from the active BES decreased after ATR RNAi (Fig.3B). Though this change was only detected as significant by RNAseq analysis after 36 hrs of RNAi (Table S3), reduction of VSG221-specific reads was seen consistently in each independently generated sample after 24 hrs of RNAi (Fig.3B) and became more pronounced from 24 to 36 hrs (Fig.3B, Fig.3 – supplementary figure 1), consistent with accumulation of cells lacking a VSG221 coat (see below). Second, the levels of increased gene-specific reads were not uniform across the silent BES but, instead, were most pronounced proximal to the promoter and telomere. Increased ESAG expression (Fig.3B, Fig.3 – supplementary figures 1 and 2) after TbATR RNAi was mainly accounted for by increased transcript abundance of the two promoter-proximal genes encoding the *T. brucei* transferrin receptor, ESAG6 and ESAG7 (Steverding et al., 1994). In addition, increased levels of transcripts were also detected after RNAi from folate transporter genes (Fig.3 – supplementary figures 1 and 2), encoded by ESAG10, which have been described downstream of a duplicated BES promoter at some telomeres (Gottesdiener, 1994; Hutchinson et al., 2016). In the silent BES the extent of increased ESAG6 and ESAG7 mapped reads after TbATR RNAi appeared broadly comparable with the extent of increased reads that mapped to the silent VSGs, suggesting similar changes at the promoter and telomere. This pattern differed from the active BES (Fig.3B, Fig.3 – supplementary figure 1, Table S3): here, the extent of reduction in reads mapping to either VSG221 or the ESAGs appeared greatest as the genes became more proximal to the telomere, whereas ESAG6 and ESAG7-specific reads appeared to increase, as seen in the silent BES. Thus, TbATR loss resulted in different transcription effects at the two ends of the active BES, distinguishing it from the silent BES.

### ATR RNAi leads to changes in VSG coat composition

To ask whether changes in VSG RNA after TbATR loss extend to VSG protein expression on the cell surface, we next performed co-IF on unpermeabilised cells before and after TbATR RNAi with antiserum recognising either VSG221 (active BES1, Telo40) or VSG121 (silent BES3, Telo4), scoring for expression of the two VSGs on individual cells (Fig.4A,B; Fig.4 – supplementary figure 1). In conjunction, flow cytometry was used to analyse larger numbers of cells, also distinguishing cells that expressed one, both or neither of the VSGs (Fig.4C; Fig.4 – supplementary figure 1). Both approaches gave comparable results, as did comparison of the two RNAi clones. In the absence of TbATR RNAi induction, >98% of cells expressed only VSG221, reflecting monoallelic control of BES transcription. RNAi led to a progressive decrease in cells that stained with only anti-VSG221 antiserum (∼80% and ∼70% of cells after 48 hrs in clones 1 and 2, respectively; Fig.4A, Fig.4 – supplementary figure 1). Concomitantly, there was a progressive increase in cells that either did not stain with antiserum against either VSG (∼5-15% after 48 hrs; Fig.4A, Fig.4 – supplementary figure 1), or stained with both anti-VSG221 and -VSG121 antiserum (∼10-15% after 48 hrs; Fig.4A,B and Fig.4 – supplementary figure 1). Cells expressing VSG121 but not VSG221 after TbATR RNAi were present, but very rare (Fig.4A,B and Fig.4 – supplementary figure 1). Detection of two VSGs on the cell surface indicates loss of monoallelic BES expression. Cells without VSG221 in the coat indicate TbATR depletion can also lead to discontinued expression of the active VSG and, presumably, expression of an undetected VSG or VSGs. Both findings are consistent with the RNAseq data.

**Figure 4.**
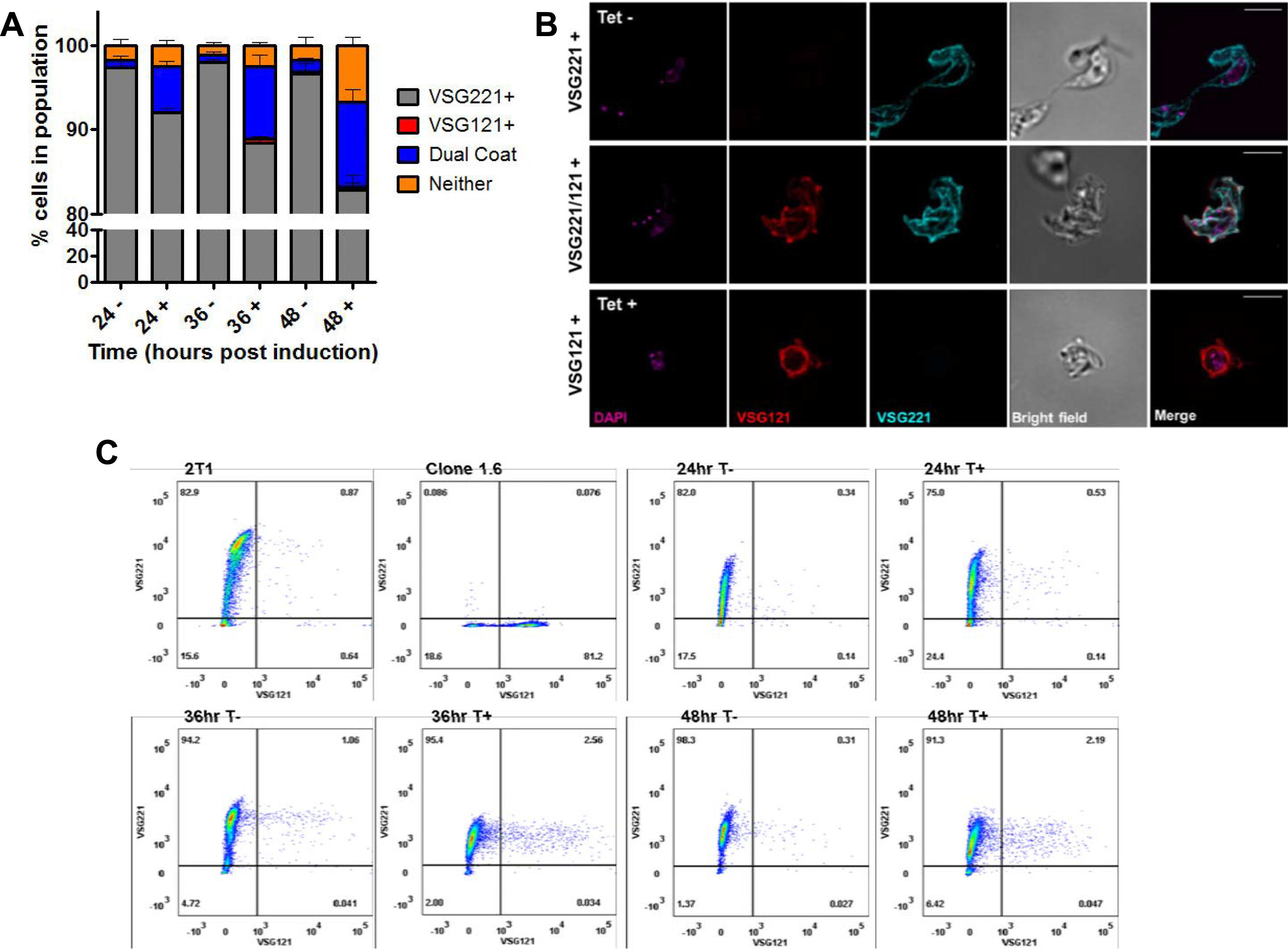
Loss of ATR in bloodstream form *T. brucei* results in changes in VSG coat expression. (A) Analysis of VSG expression by indirect immunofluorescence. Cells were collected at 24, 36 and 48 hours after TbATR RNAi induction (+) in CL1, or at the same time points without RNAi induction (−), and stained with anti-VSG221 and anti-VSG121 antiserum. Individual cells were scored for the presence of just one of the VSGs (VSG221+, grey; VSG121+, red), both VSGs (dual coat, blue), or neither VSG on their surface (orange); numbers are expressed as a percentage of the total population and error bars show ± SEM for three experimental repeats (>200 cells were counted at each time point in all experiments). (B) Representative images of CL1 cells stained with anti-VSG221 and anti-VSG121 antiserum; scale bars, 5 μm. (C) Analysis of VSG expression by flow cytometry. Cells in which TbATR RNAi had been induced (T+) or controls without induction (T−) were collected from CL1 after 24, 36 and 48 hours growth, stained with anti-VSG221 and anti-VSG121 antiserum and analysed by flow cytometry. Over 10,000 cells were analysed per sample and time point; representative date from one experiment is shown. For comparison the same analysis is shown using the parental RNAi cell, 2T1, and a cell (clone 1.6) in which expression of VSG221 has been blocked, leading to a majority of cells expressing VSG121.

### Loss of ATR leads to altered localisation of VEX1

The BES and VSG expression changes described above after TbATR RNAi display a striking overlap with those seen after RNAi against VEX1 (Glover et al., 2016; Hutchinson et al., 2016), a factor that localises specifically to the active BES and may be a component of the extranucleolar *T. brucei* ESB. To ask if the effects of loss of TbATR may be mediated through VEX1, we expressed a 12 myc-tagged variant of the factor from its own locus (Fig.5 – supplementary figure 1A)(Glover et al., 2016) in the TbATR RNAi cells, and examined expression and localisation, with and without RNAi, using anti-myc antiserum (Fig.5; Fig.5 – supplementary figure 1B, C; Fig.5 – supplementary figures 3 and 4). Cells lacking the 12myc tagged VEX1 were used as negative controls, and no staining could be seen (Fig.5 – supplementary figure 2). RNAi-mediated loss of TbATR had no discernible effect on the abundance of VEX1-12myc protein (Fig.5 – supplementary figure 1B, C), but did affect subnuclear localisation (Fig.5). Around 80% of uninduced G1-early S phase cells had a single discrete subnuclear anti-myc focus that was close to but distinct from the nucleolus, whereas two anti-myc foci were more commonly detected in S and G2 phase cells. Most post-mitotic cells had two anti-myc foci: one in each nucleus. These distributions compare well with VEX1-12myc analysis described by Glover et al, 2016, and the small numbers of cells with three or more anti-myc foci, which were not described previously (Glover et al., 2016), may be explained by ‘leaky’ RNAi in the absence of tetracycline induction. TbATR RNAi substantially changed the distribution of anti-myc signal (Fig.5A,B). First, in all cells that could be assigned a clear cell cycle stage, increased numbers of cells with two and (in particular) three or more anti-myc foci were seen (examples shown in Figs. 5B, Fig.5 – supplementary figures 3 and 4). Second, cells were readily detected where the anti-myc signal overlapped with the nucleolus, an arrangement that was barely detected in uninduced cells. Finally, in the aberrant cells generated by TbATR RNAi, which do not conform to expected nuclear and kinetoplast DNA cell cycle configurations, virtually none had a single anti-myc focus; instead, most had three or more foci, and overlap of anti-myc signal with the nucleolus was mostly readily detected here. Taken together, these data reveal that loss of ATR in BSF *T. brucei* perturbs the localisation of VEX1, causing the protein to be found in more than one location, and losing its clear spatial separation from the nucleolus.

**Figure 5.**
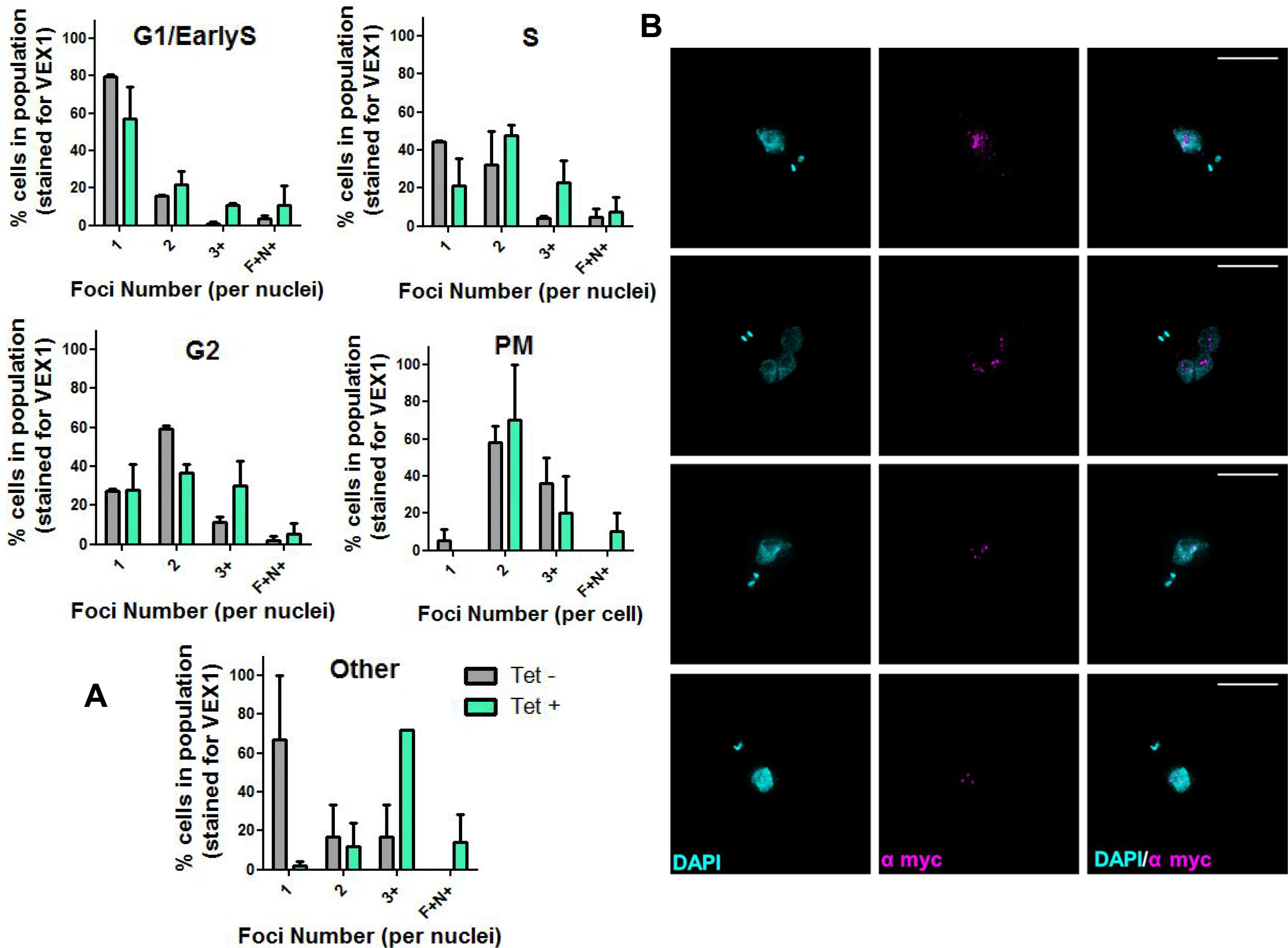
Altered localisation of VEX1 after ATR RNAi. (A) Analysis of VEX1 foci number and localisation by indirect immunofluorescence. Cells were collected after 24 hours growth with (Tet+, cyan bars) or without (TeT−, grey bars) induction of TbATR RNAi and stained with anti-myc antiserum to detect VEX1-12myc. Nuclear (N) and kinetoplast (N) DNA was stained with DAPI and used to categorise individual cells into G1/Early S, S, G2 and posT−mitosis (PM) stages of the cell cycle; cells that deviated from expected N-K configurations and were scored as ‘other’. For each cell cycle stage (and for other cells) the presence of 1, 2, 3+ (3 or more) anti-myc foci that were distinct from the nucleolus, or where the anti-myc focus (or foci) and nucleolus overlapped (F+N+), were scored, and their numbers are expressed as a percentage of the total number of cells counted. Errors bars represent SEM from three experimental replicates. (B) Representative images of VEX1-12myc localisation observed in cells grown for 24 hours of TbATR RNAi induction; DNA is stained with DAPI (cyan) and anti-myc antiserum (magenta) was used to detect VEX1-12myc. Images were captured on an Elyra super resolution microscope; scale bar = 5 µm.

### Loss of ATR alters localisation of RNA Polymerase I

Since *VSG* transcription is catalysed by RNA Pol I sequestered to the actively transcribed BES, we next asked if the altered VEX1 localisation after TbATR RNAi was reflected in changed RNA Pol I localisation. To do so, immunofluorescence with anti-RNA Pol I antiserum (Glover et al., 2016) was compared in cells grown for 24 hrs with or without TbATR RNAi, the earliest time point at which growth was impaired, and when VSG expression and VEX1 localisation changed (Fig. 6). Consistent with previous reports (Kerry et al., 2017; Navarro and Gull, 2001), around 55% of uninduced cells showed two anti-RNA Pol I signals (Fig.6, Fig.6 – supplementary figure 1), corresponding to a single nucleolus and ESB. Though TbATR RNAi resulted in only a modest increase (from ∼45% to ∼55%) in the numbers of cells lacking two separate RNA Pol I signals (Fig.6 – supplementary figure 1), this appeared consistent with the increased numbers of cells in which VEX1-12myc overlapped with the nucleolus (Fig.5A). The more striking effect of TbATR RNAi was an increase in the number of cells with more than two anti-RNA Pol I subnuclear foci (Fig.6B). It seems likely, therefore, that the increased numbers of RNA Pol I foci correspond with the increased numbers of VEX1-12myc foci after loss of TbATR, suggesting the subnuclear perturbations are connected.

**Figure 6.**
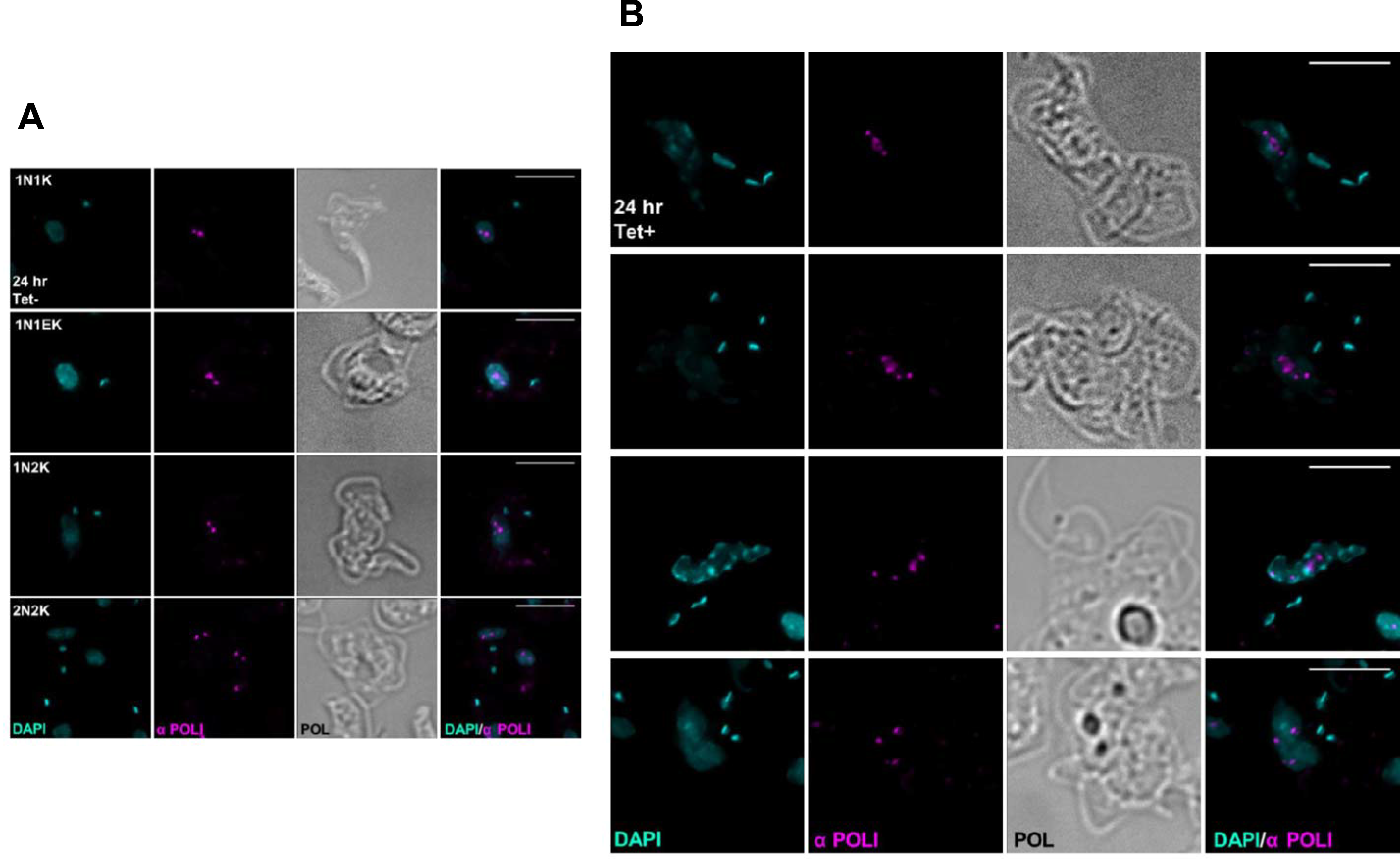
Altered localisation of RNA Polymerase (pol) I after depletion of ATR by RNAi. (A) Representative images of *T. brucei* bloodstream form cells grown in the absence of TbATR RNAi (24 hrs, TeT−)stained with anti-RNA Pol I antiserum (magenta); cells are shown in the different stages of the cell cycle, determined by configurations of nuclear (N) and kinetoplast (K) DNA stained with DAPI (cyan). (B) Representative images of RNA Pol I localisation in cells grown for 24 hours with induction of TbATR RNAi (Tet+). Images were captured on a DeltaVision microscope; scale bars = 5 µm.

**Figure 7.**
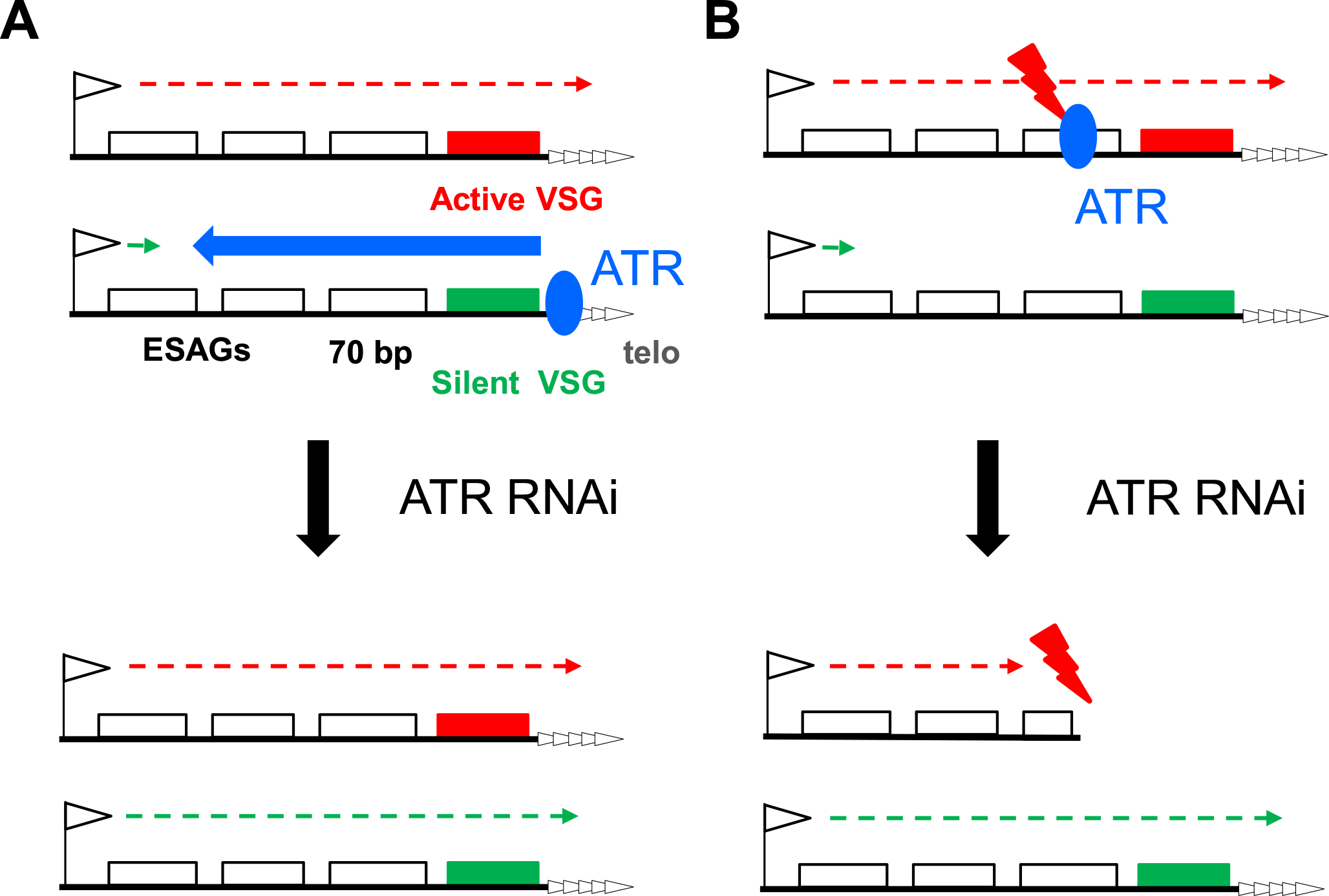
Two models for the role of ATR in controlling *T. brucei* surface antigen gene expression. (A) Transcription (green arrow) from the RNA Pol I promoter (arrow) of a silent BES is suppressed (blue arrow) by ATR (blue circle) and does not traverse the ESAGs (white boxes), 70 bp repeats (hatched box), VSG (green box) or telomere repeats (arrayed arrow heads). In contrast, TbATR does not impede transcription (red arrow) of the single active BES (active VSG, red). After TbATR RNAi, silencing is compromised and transcription can extend across the silent BES, leading to expression of the normally silent VSG. (B) TbATR recognises and signals the repair of lesions (red lightning) within the actively transcribed VSG BES. Loss of TbATR by RNAi means lesions are not effectively repaired and integrity of the active BES is compromised (e.g. loss of VSG), which is lethal and selects for cells expressing a silent VSG BES.

## Discussion

In this work we provide functional analysis of the damage signalling protein kinase ATR in the eukaryotic parasite *T. brucei*. Our data reveal that loss of TbATR is severely detrimental to mammal-infective parasites, with RNAi resulting in increased nuclear damage and genotoxin sensitivity. In addition, we show that TbATR loss impairs the control mechanism(s) adopted by *T. brucei* the ensure monoallelic expression of its critical VSG coat, including the subcellular localisation of the BES control factor VEX1 and RNA Pol I. Together these data indicate conserved roles for *T. brucei* ATR in recognising and reacting to lesions or perturbations that affect nuclear genome function, as well as parasite-specific functions in maintaining gene expression needed for immune evasion and transmission.

Cessation of BSF *T. brucei* growth after RNAi suggests TbATR is essential in BSF cells, consistent with previous kinome-wide growth analysis (Jones et al., 2014) and RITseq (Fernandez-Cortes et al., 2017; Stortz et al., 2017). Such critical ATR importance is also found in yeast (Cha and Kleckner, 2002) and during mammal development (Brown and Baltimore, 2000), but is not universal across eukaryotes, since ATR null mutants are viable in plants (Culligan et al., 2004). What TbATR functions are essential for *T. brucei* are unclear. One possibility is a role for TbATR in a critical DNA repair pathway, since loss of the PK results in increased sensitivity to damage caused by gamma irradiation, alkylation (MMS), nucleotide depletion (HU) and DNA cross-links (UV), indicating TbATR plays roles in co-ordinating the response to a range of genotoxic lesions. Such roles are also consistent with elevated levels of γH2A and focal accumulation of RAD51 after RNAi, similar to effects observed after ATR-depletion in human cells [1] and indicating loss of *T. brucei* ATR results in increased levels of endogenous nuclear genome damage. γH2A expression in *T. brucei* and *L. major* has been shown to increase after induction of a DSB or exposure to MMS, phleomycin or HU (Damasceno et al., 2016; Glover and Horn, 2012), indicating phosphorylation occurs due to a range of lesions and may not simply reflect DSB formation (Revet et al., 2011; Turinetto and Giachino, 2015).

Similarly, though focal accumulation of *T. brucei* RAD51 after TbATR loss may reflect association of the recombinase with single-stranded DNA, it is unclear whether increased foci arise due to increased abundance of processed DSBs, replication damage, aberrant telomeres or other lesions. Indeed, it was notable that accumulation of RAD51 foci was slower than increased γH2A expression after TbATR RNAi, perhaps indicating the lesions that first result from loss of the PIKK are not bound by RAD51.

Amongst the range of potential nuclear genome lesions that *T. brucei* ATR might act on, DSBs have been detected within the BES (Boothroyd et al., 2009; Glover et al., 2013a) and their targeted generation by ISceI endonuclease impairs *T. brucei* BSF survival (Devlin et al., 2016b; Glover et al., 2013a; Glover et al., 2011). However, whether DSBs reach endogenous levels that are a threat to *T. brucei* viability without TbATR is unclear, in particular because kinome-wide RNAi analysis suggests TbATM (which is predicted to recognise DSBs directly) is not essential (Jones et al., 2014; Stortz et al., 2017). In other eukaryotes ATR has a key role in recognising and responding to replication stress (Yazinski and Zou, 2016), including regulating the timing of replication origin firing and activation of back-up (dormant) origins (Chen et al., 2015; Shechter et al., 2004), stabilisation and protection of stalled replication forks (Dungrawala et al., 2015; Hu et al., 2012), and replication at fragile sites (Barlow et al., 2013; Casper et al., 2002). It is conceivable that near genome-wide multigenic transcription in *T. brucei* and other kinetoplastids results in pronounced replication stress due to localised, frequent clashes between DNA and RNA Pol complexes; indeed, it has been argued that a need to limit such collisions may have selected for widely-spaced, rigidly positioned replication origins in *T. brucei* (Devlin et al., 2016b; Tiengwe et al., 2012). However, mapping such putative clashes is needed to determine if TbATR contributes to their resolution, as is evaluating whether the PK’s role and interaction partners in the response are conserved or diverged. HU-derived replication stress leads to increased activity of one origin in chromosome 1 of *T. brucei* (Calderano et al., 2015), but HU effects on replication have not been mapped genome-wide or in the absence of TbATR. Since highly unconventional replication responses to HU stress have been described in *Tetrahymena* (Sandoval et al., 2015), the *Leishmania* 9-1-1 complex has been shown to provide essential, non-canonical replication-associated functions (Damasceno et al., 2016), and origin designation and replication progression may be divergent in both *Leishmania* and *T. brucei* (Marques et al., 2015; Marques et al., 2016), it is not clear that characterised ATR replication roles can be simply extrapolated to kinetoplastids, or other eukaryotic microbes. Nonetheless, ATR in other eukaryotes has also been implicated in the repair of DNA damage, including crosslinks, acted upon by nucleotide excision repair (NER)(Munoz et al., 2017; Ray et al., 2013) and, intriguingly, the NER pathway in *T. brucei* appears to have diverged in its operation (Badjatia et al., 2013; Machado et al., 2014), perhaps as an essential adaptation to multigenic transcription. In addition, a nuclear translesion DNA Pol has been shown to be essential (Rudd et al., 2013), with its loss stalling DNA replication. Thus, TbATR’s potential roles in responding to replication stress appear worthy of further investigation, since divergence due to the demands of multigenic transcription may be an attractive drug target, such as by anti-cancer drugs that target ATR (Yazinski and Zou, 2016).

Monoallelic BES expression leading to a single VSG coat on the surface of a BSF *T. brucei* cell is central to the success of immune evasion by antigenic variation, with parallel processes used in many pathogens (Glover et al., 2013b; Guizetti and Scherf, 2013; Obado et al., 2016). Precisely how a single BES is selectively transcribed while the remaining ∼15 are largely transcriptionally silent is still being unravelled (Gunzl et al., 2015). The range of factors identified whose loss undermines monoallelic expression encompasses chromatin modifiers, chromatid cohesin proteins, telomere-associated proteins, nuclear lamina components, inositol phosphate signalling and, recently, a novel ESB-associated factor, VEX1 (see introduction for specific references). To date, DNA repair factors have not been strongly implicated in monoallelic expression, but instead with VSG switching by recombination. However, it might be noted that mutation of the HR factors RAD51 (McCulloch and Barry, 1999), BRCA2 (Hartley and McCulloch, 2008) and RAD51-3 suppresses levels of antigenic variation not only by impairing VSG recombination, but also by lowering levels of transcriptional switching between BES. In addition, exposure of BSF *T. brucei* to DNA damaging agents can increase silent BES transcription (Sheader et al., 2004). Is it possible that the effects revealed by RNAi of TbATR reveal a closer than anticipated link between BES transcriptional control and the DNA damage response? RNAseq (validated by RT−qPCR of specific VSGs from silent BES and examination of the surface coat) shows that loss of TbATR has an early and profound effect on BES gene expression, leading to impairment in active BES transcription, transcription from previously silent BES and the expression of at least two VSGs on the cell surface. Remarkably, the effects we describe display considerable overlap with those seen after RNAi of *T. brucei* VEX1, a factor that associates with the active BES in BSF cells (Glover et al., 2016). Like the effects of TbATR RNAi, loss of VEX1 results in increased abundance of silent BES VSG RNAs, decreased expression of VSG221 protein, co-expression of at least two VSGs on the cell surface, and little evidence for singular expression of a previously silent VSG (VSG121/VSG6). Moreover, RNAseq mapping to the silent BES shows the same pattern of increased transcripts after TbATR and VEX1 RNAi (Hutchinson et al., 2016): comparable increases in promoter- and telomere-proximal gene-specific reads, but little evidence for changed levels of ESAGs centrally located in the BES (an effect distinct from VEX1 overexpression). In only two ways do the available data for TbATR and VEX1 diverge. First, increased abundance of non-BES RNA Pol I transcripts, including procyclin and PAG, are seen in our data but were only observed after VEX1 overexpression, not knockdown (Glover et al., 2016).

This difference may reflect wider roles for TbATR in genome maintenance, or the application of different thresholds of significance when comparing RNAi induced and uninduced RNAseq data. Second, whereas loss of TbATR results in reduced RNA from the telomere-proximal VSG221, as well as from at least some ESAGs in the active BES, no such loss of transcripts was described after VEX1 RNAi. It seems plausible that this difference may be explained by lack of damage repair after TbATR RNAi, a function that may not be provided by VEX1, as discussed below. Nonetheless, the perturbations in VEX1 subnuclear localisation we describe after TbATR RNAi reinforce a connection between the PIKK damage repair kinase and this novel factor in BES expression control. Whether this connection is direct or indirect is currently unclear.

Two scenarios might be considered to explain the loss of monallelic VSG expression after TbATR RNAi and the overlap in phenotypes seen after VEX1 RNAi (Fig.6). One explanation is that TbATR plays an active role in exerting monoallelic transcriptional control on the BES (Fig.6A). One route for such a function could be through TbATR interaction with telomeres, which resemble DSBs and are therefore protected by the shelterin complex to avoid eliciting a damage response (Feuerhahn et al., 2015). In other eukaryotes one shelterin component, POT1, interacts with ATR to prevent its activation (Denchi and de Lange, 2007), while ATR (and ATM) have been shown to recruit telomerase (Moser et al., 2011; Tong et al., 2015) and, perhaps, shelterin (Moser et al., 2009) to telomeres. Shelterin binding extends to subtelomeres and can cause silencing of genes through the telomere position effect (TPE)(Ottaviani et al., 2008). In *T. brucei* RNAi of RAP1 (Yang et al., 2009), TRF (Jehi et al., 2014a) and TIF2 (Jehi et al., 2014b), each a shelterin factor, results in impaired BES silencing or VSG switching, suggesting some parallels with the effects of TbATR loss described here. However, a complication is that measurements of TPE in *T. brucei* indicate silencing stretches for only a few kb (Glover and Horn, 2006) and does not encompass the whole BES, which seems at odds with increased promoter-proximal transcripts levels after TbATR or VEX1 loss. In addition, it seems unlikely that telomere integrity is the basis for such an activity, since, in contrast to the rapid BES transcription changes seen after TbATR RNAi, telomere repeat attrition after mutation of telomerase is slow to accumulate (Dreesen et al., 2005) and excision of the telomere tract in active or silent BES does not elicit a change in BES transcription or switching (Glover et al., 2007; Glover et al., 2013a). Nonetheless, it is conceivable that TbATR, like VEX1, acts to exert transcriptional controls at both the BES telomere and promoter. Alternatively, TbATR may not act directly on the BES, but its loss might impair integrity of the nucleolus, perhaps because the kinase acts to sense nucleolar structure (Kidiyoor et al., 2016) or responds to DNA damage that impedes RNA Pol I transcription (Larsen and Stucki, 2016). Such an effect may explain increased RNA Pol I foci after TbATR RNAi, leading to increased VEX1 foci. However, this explanation would imply the ESB, and perhaps VEX1 activity, is an extension of the nucleolus. Though such a connection appears consistent with observations that inhibition of RNA Pol I transcription leads to apparently concurrent breakdown of the nucleolus and loss of extranucleolar RNA Pol I and VEX1 signal (Kerry et al., 2017), it is at odds with evidence for the ESB being a discrete subnuclear structure (Navarro and Gull, 2001). Thus, given the alterations we see in VEX1 localisation after TbATR RNAi, it will be valuable to determine if TbATR and VEX1 act together to influence the deposition or activity of related factors at the BES (active and silent). However, currently no activity has been ascribed to VEX1 and whether TbATR displays any localisation to the BES is complicated by its widespread distribution throughout the nucleus (data not shown), reflecting wider roles in genome maintenance. Nonetheless, it is possible that VEX1 is phosphorylated by TbATR, either directly or via a signalling network, altering its activity. In this regard, RNAseq suggests significantly reduced levels of VEX1 RNA 36 hrs after TbATR RNAi (data not shown), though this may be a secondary result of widespread impediment to gene expression due to unrepaired damage.

A radically different explanation for the effects we describe after RNAi is that loss of TbATR affects integrity of the active BES (Fig.6B), consistent with ATR’s well-established role in signalling and co-ordinating the repair of DNA lesions. Recent work has begun to test for lesions that might initiate VSG switching, leading to proposals for generation of DSBs within the active BES (Boothroyd et al., 2009; Glover et al., 2013a), telomere fragility as a source of subtelomeric DNA breaks (Hovel-Miner et al., 2012; Jehi et al., 2014b), and damage arising from early DNA replication of the active BES (Devlin et al., 2016a; Devlin et al., 2016b).

TbATR could conceivably recognise and signal any such VSG switch-initiating lesion and absence of the PIKK after RNAi might then provide an explanation for the reduced telomere-proximal gene-specific transcripts: unrepaired lesions could lead to loss of sequences from the active BES, perhaps because of a failure to halt cell cycle progression to allow repair. Such an explanation would explain the difference in RNAseq mapping at the telomere of the active BES in TbATR and VEX1 RNAi cells (assuming VEX1 has no repair functions).

Moreover, the apparent increasing loss of gene-specific RNAs with greater proximity to the telomere may indicate there is no single site of lesion generation, but instead increasing damage from promoter to telomere. In this regard, the established role of ATR in tackling transcription-replication clashes is intriguing (Hamperl et al., 2017), since putative lesion density would follow the direction of transcription, explaining the pattern of gene-specific transcript loss after TbATR RNAi in the active BES. Moreover, the actively transcribed BES is replicated earlier in S-phase than the silent BES (Devlin et al., 2016b). Thus, if damage in the active BES was not signalled by TbATR, leading to switching or loss of monoallelic expression (or both), the timing of BES replication may breakdown, with greater than a single BES replicated early in S-phase, causing each to be bound by VEX1. Chromosome miss-segregation due to loss of TbATR may have a similar effect, resulting in more than a single VEX1 focus in divided cells. However, this model does not readily explain three outcomes of TbATR RNAi: increased RNA from promoter-proximal genes in the silent BES, the expression of at least two VSGs on the parasite surface, and loss of VEX1 spatial separation form the nuclelous. The former outcome may be explained as a stress response to reduced transferrin receptor expression (perhaps due to loss of integrity of the active BES), as has been described previously (van Luenen et al., 2005). However, why this outcome is also seen after VEX1 RNAi is less clear. Though detection of cells lacking VSG221 expression indicates TbATR RNAi can lead to loss of the active VSG, the infrequent detection of RNAi induced cells expressing just VSG121 and the ready detection of cells expressing two VSGs indicates VSG switches by recombination are not the sole consequence of TbATR knockdown. As a result, loss of damage sensing by TbATR also leads to loss of BES monallelic expression, linking DNA damage repair and transcriptional control. Indeed, such a link may explain previously detected events in which transcriptional VSG switching and deletion of the active BES occur in concert (Cross et al., 1998; Rudenko et al., 1998), perhaps indicating a common initiating event for the two strategies of antigenic variation used by *T. brucei*. VEX1 co-localisation with the nucleolus after TbATR RNAi may suggest that ATR does not simply tackle BES DNA damage, but also plays roles in monitoring integrity of the nucleolus (as discussed above). Characterising the nature of the lesions TbATR acts upon, the signalling targets of TbATR, and the roles of putatively ESB-associated factors such as VEX1 will test these possibilities.

## Materials and methods

### Cells and RNAi analysis

To examine the role of ATR in BSF *T.brucei*, a previously described construct containing an RNAi target sequence derived from the coding sequence of TbATR was generated using the Gateway cloning strategy as described by(Jones et al., 2014). The construct was transformed into the 2T1 parental cell line and two clones were recovered for further analysis. Cells were grown in HMI-9 medium supplemented with 20 % (v/v) foetal calf serum (Stortz et al., 2017) and RNAi cells were maintained in the 5 µg.ml^−1^ Hygromycin and 5 µg.ml^−1^ Phleomycin. For maintaining myc tagged expressing cell lines, 10 µg.ml^−1^ Blasticidin was added to the media. To assess RNAi knockdown, one allele of the TbATR gene was endogenously tagged at the C-terminus with 12 copies of the myc epitope (12myc) using the vector pNATx12myc (kind gift, D. Horn). Cloning was conducted as described in (Devlin et al., 2016b). Growth curves were performed as described in (Stortz et al., 2017). For those performed in the presence of genotoxic stress, the following concentration or exposure of genotoxic agents were used (unless stated otherwise): MMS (0.0003%), hydroxyurea (0.06 mM), UV (1500 J/m^2^) and IR (150 Gy). For UV and IR, RNAi was induced for 24 hours prior to exposure. To assess gene knockdown using RT−qPCR, RNA was extracted using the RNeasy kit (Qiagen manufacturer’s instructions). RNA was treated for 30 minutes at room temperature off column with DNase (Qiagen) to minimise DNA contamination. cDNA was prepared as per manufacturer’s instructions using Superscript III^®^ Reverse Transcriptase (RT; Thermo Fischer) using random hexamers. RT minus samples were prepared to control for genomic DNA contamination.

Oligonucleotide sequences were designed using Primer Express^®^ 3.0 software (Applied Biosystems). The following master mix was set up: 2.5 µl of the appropriate cDNA, 2.5 µl of the appropriate primers (300 nM stock) and 12.5 µl SYBR^®^ Green PCR Master Mix (Applied Biosystems) in a total volume of 25 µl. Samples were set up in a MicroAmp^®^ Optical 96-well Reaction Plates (Thermo Fischer) as triplicates and run in a 7500 Real Time PCR system (Applied Biosystems). The following PCR conditions were used: 50 °C for 2 min (x 1), 95 °C for 10 min (x 1), 95 °C for 15 sec followed by 60 °C for 1 min (x 40) followed by a *dissociation step (95 °C for 15 secs, 60 °C for 1 min, 95 °Cfor 15 secs and finally 60 °C for 15 secs). Data* was analysed using the ΔΔCt method (Schmittgen and Livak, 2008). Oligonucleotide sequences are available on request.

### Immunoblotting and Immunofluorescence

Immunoblotting to assess levels of γH2A, or to detect myc-tagged proteins or RAD51, was performed and analysed as described in (Stortz et al., 2017). Immunofluorescence of surface VSGs was performed as described in (Glover et al., 2016) with the following modifications: anti-VSG121 and anti-VSG221 were both used at a concentration of 1:8000; goat anti-rabbit AlexaFluor 488 or goat anti-rat AlexaFluor 594 (Invitrogen) secondary antisera were used. Immunofluorescence of VEX1-12myc and RNA Pol I was performed as described in (Glover et al., 2016).

### RNAseq analysis

Cells were sampled at 24 and 36 hrs post RNAi induction, or at equivalent times without induction, from two biological replicates of TbATR CL1 cells and from a single replicate of TbATR CL2, providing triplicate induced and uninduced samples at both time points. In all cases cells were harvested and RNA prepared as described above. PCR analysis was performed on the isolated RNA to test for the presence of genomic DNA contamination; no contamination could be detected (data not shown). RNA concentration was measured using a Qubit (Thermo Fischer) as per the manufacturer’s instructions. An RNAseq library was prepared for each sample using a TruSeq Stranded mRNA Library Prep Kit (Illumina). The libraries were paired-end sequenced using an Illumina NextSeq 500 and a Mid-Output Flow cell, generating read lengths of 75 bp. The total number of reads for each sample, the percentage mapped and the number and percentage properly paired were assessed for each sample and found to be comparable (data not shown). To assess the distribution of sequence fragments per kilobase transcript per million bases (fpkm) across each individual replicate, a box plot was generated in RStudio using the ‘csboxplot’ command, plotting the log10 transformed fpkm for each read. The box plots for each individual replicate at 24 and 36 hrs, with or without RNAi induction, showed the spread of data for all replicates at both time points to be comparable, with similar interquartile ranges of ∼ 1 (for 24 hrs) and 1.25 (36 hrs) and medians of ∼2 (for all replicates; data not shown). To ask what transcripts displayed altered levels after TbATR RNAi, fold change (log2 transformed FC) in average fpkm for the three replicates was plotted on a volcano plot relative to the significance for each gene (the log10 transformed p-value), using the CuffDiff software in RStudio; the data was plotted using ggplot2. RNAseq reads were aligned to the Lister 427 VSG ES (Hertz-Fowler et al., 2008) using HISAT2 (Pertea et al., 2016) in ‘no splice alignment’ mode; reads with a MapQ value <1 were removed using SAMtools, which has been shown to remove >99% of short read alignment to the wrong ES (Hutchinson et al., 2016). Read mapping was visualised using Matplotlib and a custom python script.

### Analysis of VSG expression

Flow cytometry was performed as described in (Glover et al., 2016) to identify VSG121 or VSG221 positive cells: signal from excitation with the BB515 laser (BB515, log) was plotted against the signal from the excitation with the PE-CF594 laser (PE-CF594, log), using as controls 2T1 cells that predominantly express VSG221 and clone 1.6 cells (Glover et al., 2007) that predominantly express VSG121.

### Imaging and image processing

Images captured on an Axioskop2 (Zeiss) fluorescent microscope used a 63x lens and images were acquired with ZEN software. For images captured on an Olympus IX71 DeltaVision Core System (Applied Precision, GE), a 1.40/100 × lens was used and images were acquired using SoftWoRx suit 2.0 (Applied Precision, GE). Z-stacks were acquired and images de-convolved (conservative ratio; 1024×1024 resolution). Super-resolution structural illuminated images were captured on an Elyra PS.1 super resolution microscope (Zeiss), and using images were acquired using ZEN software as Z-stacks.

ImageJ (Fiji; [3]) was used to remove the background of images and for counting cells. False colours were assigned to fluorescent channels. 3D images were generated using IMARIS software (V8.2). Scale bars are as stated on images or in legends.

## Acknowledgements

We thank all current and previous members of the McCulloch and Mottram labs for input. We thank Sebastian Hutchinson for providing the reference genome files used for RNAseq analysis, and Lucy Glover and David Horn, plus lab members, for providing anti-VSG121 and anti-RNA Pol I antiserum, as well as the VEX1-12myc-tagging construct, and for discussions. This work was supported by the BBSRC [BB/K006495/1, BB/M028909/1, BB/N016165/1 and a DTP studentship to J.A. Black]. The Wellcome Centre for Molecular Parasitology is supported by core funding from the Wellcome Trust [104111]

## Data Access

Sequences used in the mapping have been deposited in the European Nucleotide Archive (accession number PRJEB23973).

## Figure supplements

**Fig.1 – supplementary figure 1.**
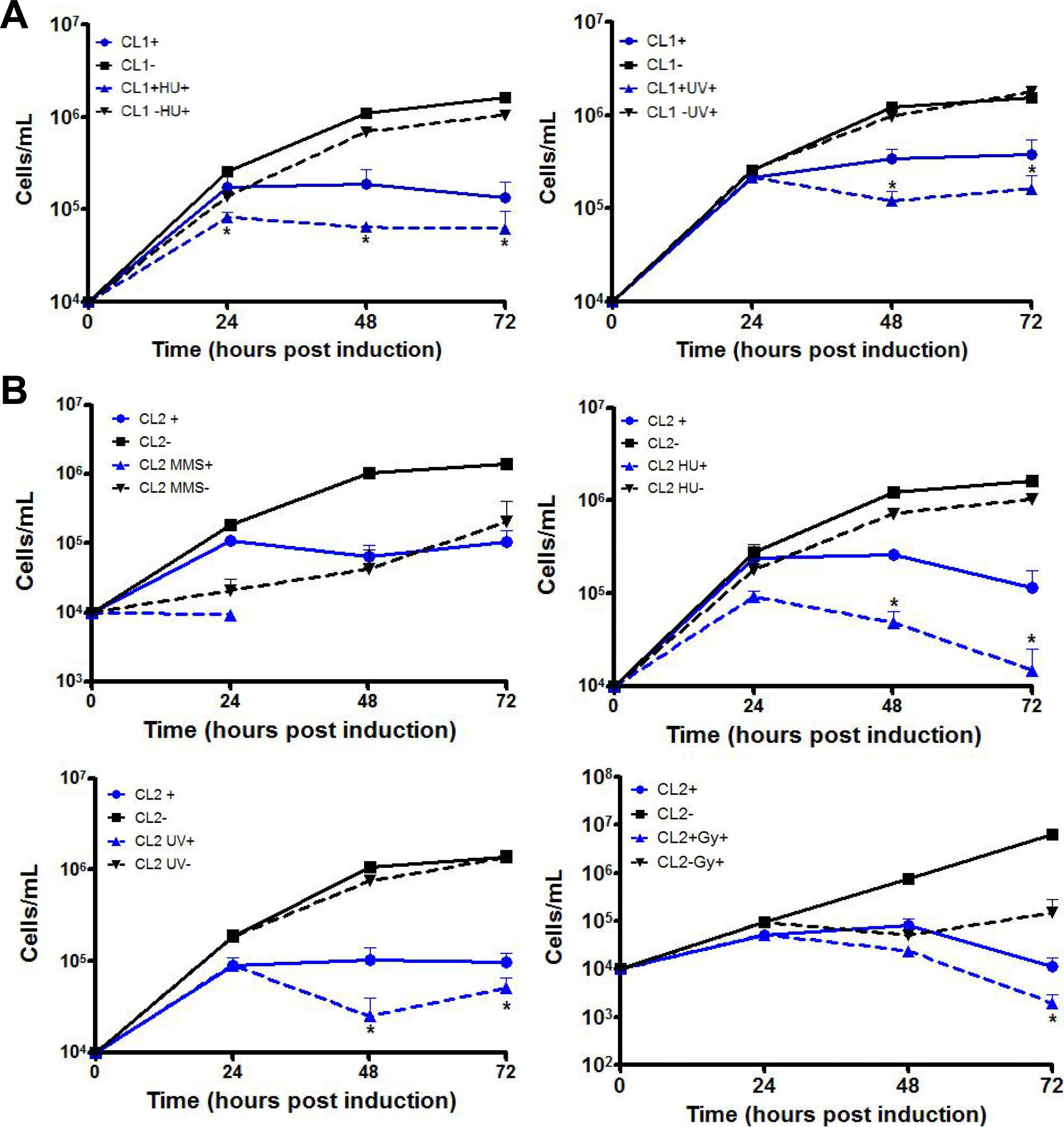
Effect of RNAi induction against TbATR on growth of BSF *T. brucei* cells in the presence of genotoxic stress. Growth curves, showing cell density with time, with (+) and without (−) induction of TbATR RNAi in the presence of a variety of genotoxic stress sources: MMS (0.0003%), UV (1500 J/m^2^), HU (0.06 mM) and IR (150 Gy). (A) Data after HU and UV treatment for clone 1 (CL1). (B) Data after all treatments shown for clone 2 (CL2). In all cases error bars show ± SEM from three experimental replicates.

**Fig.1 – supplementary figure 2.**
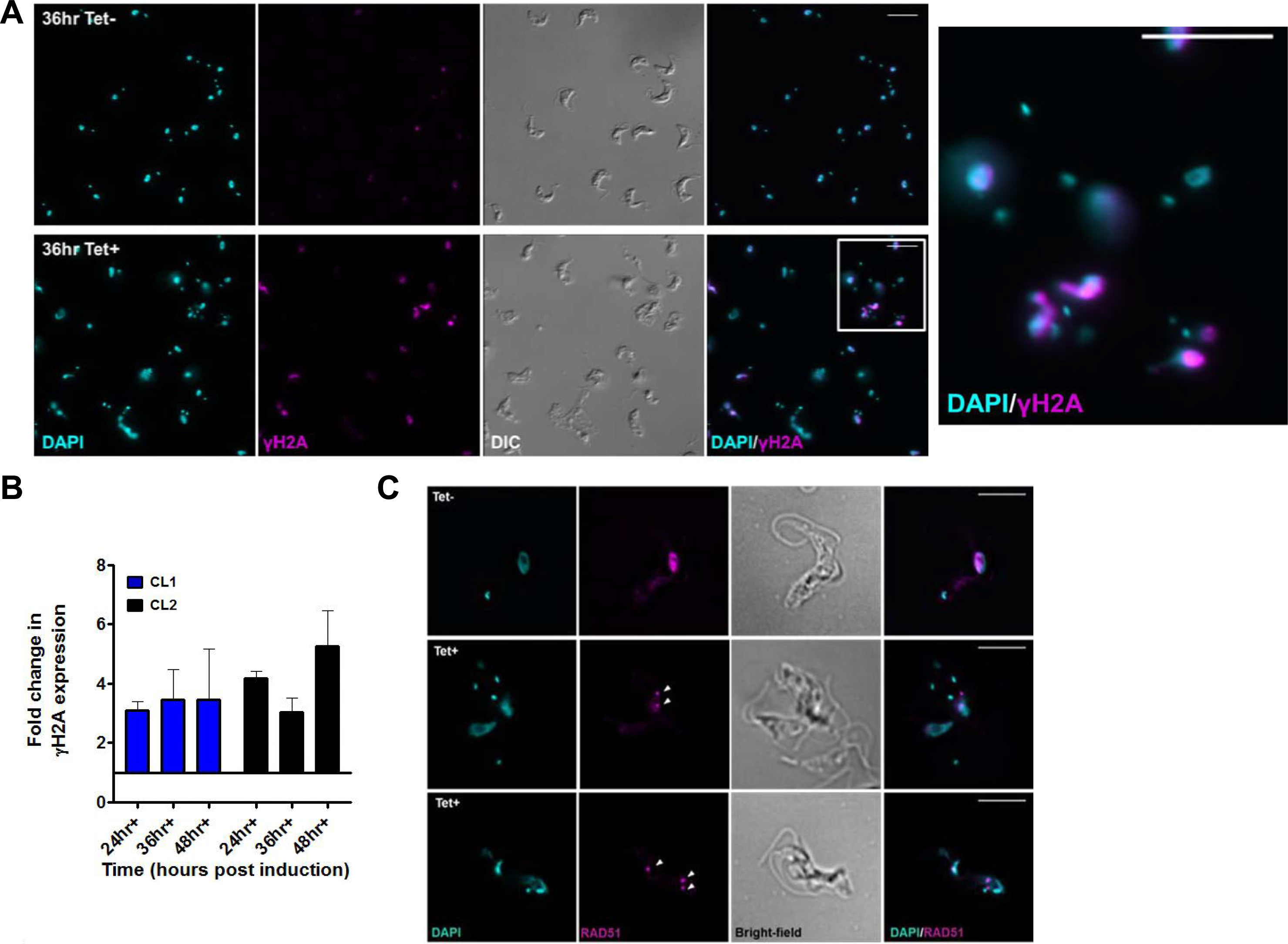
Nuclear DNA damage after depletion of TbATR by RNAi in BSF cells. (A) Cells were collected 36 hours after induction of TbATR RNAi (Tet+), or without induction (TeT−), and stained with anti-γH2A antiserum (magenta). DNA was stained with DAPI (cyan). Images (from clone 1; CL1) are shown of a relatively wide field to capture a large number of cells; scale bar=10 μm. An enlarged image of RNAi induced cells, highlighted by a white box, is shown at higher magnification. (B) Fold-change in levels of γH2A protein in clones CL1 and CL2 after 24, 36 or 48 hrs growth with (+) RNAi induction; values are shown relative to uninduced cells (set at 1) after normalisation using EF1α signal. (C) Representative images of RAD51 localisation by indirect immunofluorescence with anti-RAD51 antiserum (magenta) after 36 hrs growth with (+) or without (−) RNAi induction; DNA is shown stained with DAPI (cyan), a merge of the RAD51 and DAPI signals is provided, and cell outline in shown as bright field images (Scale bar = 5 μm; arrows highlight RAD51 foci).

**Fig.3 – supplementary figure 1.**
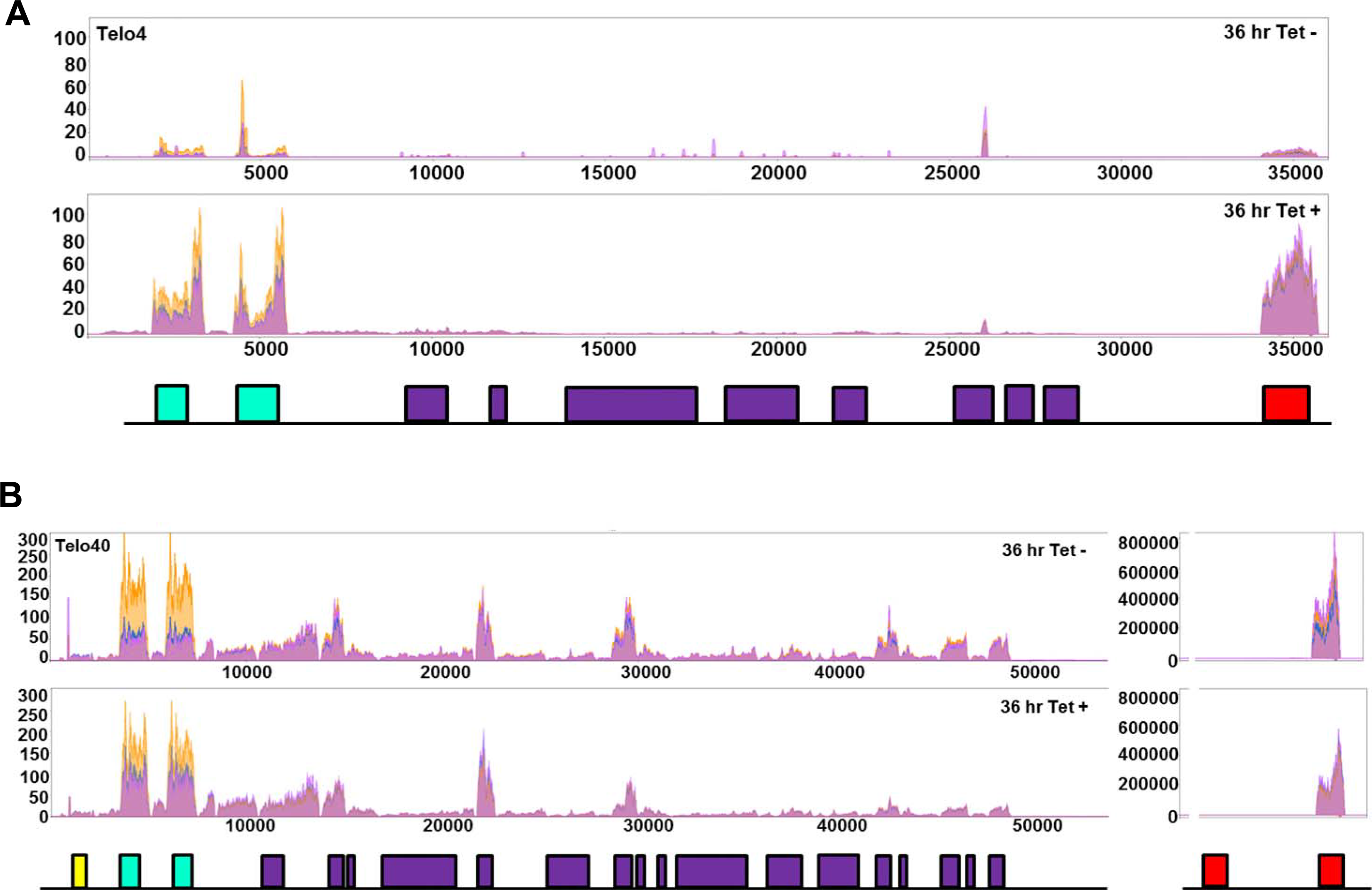
Loss of ATR impairs control of *T. brucei* bloodstream form VSG expression site transcription. Mapping of MapQ filtered RNAseq reads is shown across a silent VSG bloodstream VSG expression sites (Telo4) and the single active site (Telo40) after 36 hrs growth with (T+) or without (T−) induction of TbATR RNAi; y axes show the number of reads that map relative to location in the transcription units, and data from three replicates are overlaid. ESAG6 and ESAG7 genes are shown in green, all other ESAGs are in purple, the VSG is in red, and a gene encoding resistance to hygromycin is in yellow. For Telo40, read mapping to the VSG is shown on a different scale to the rest of the expression site.

**Fig.3 – supplementary figure 2.**
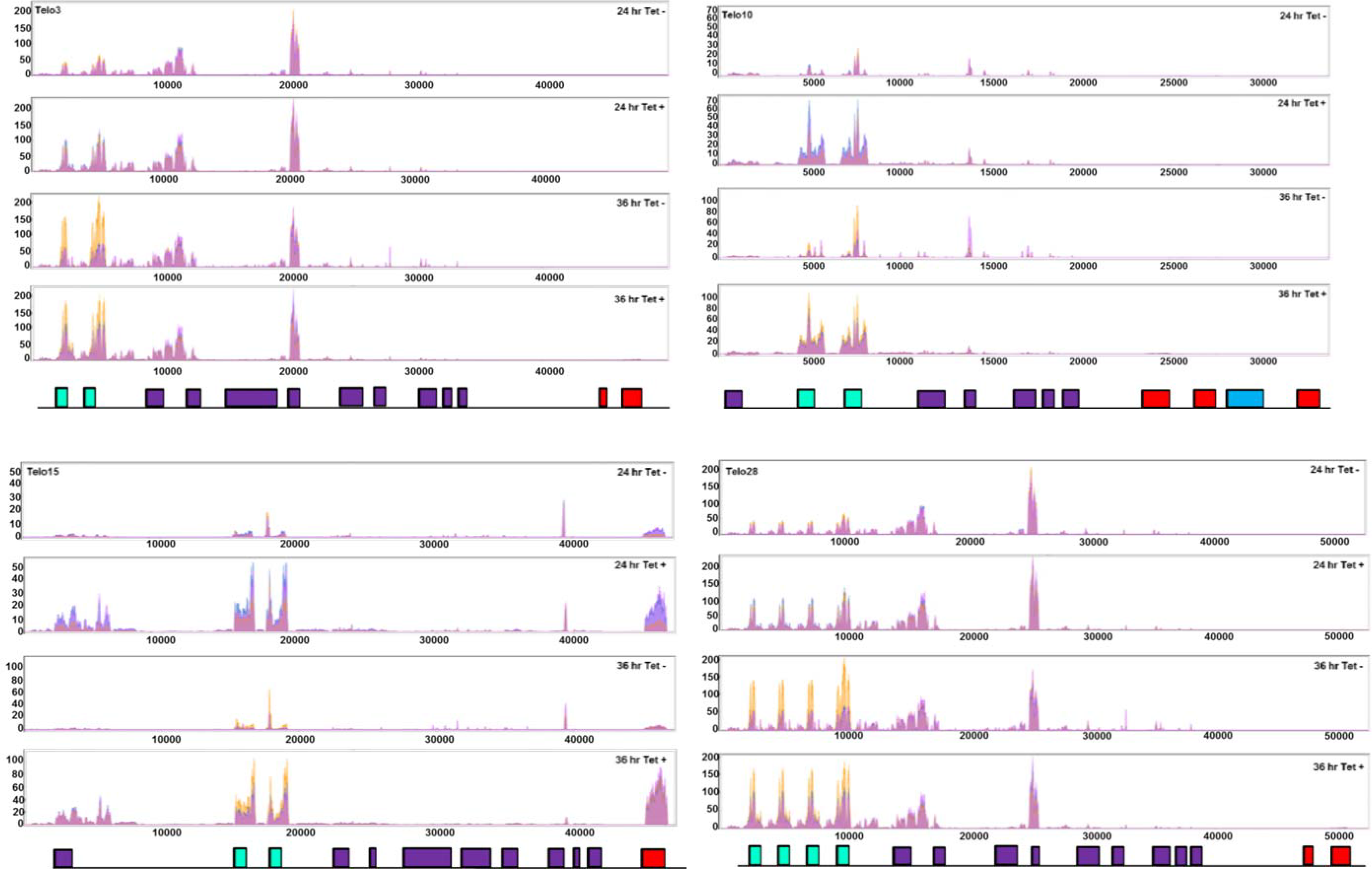
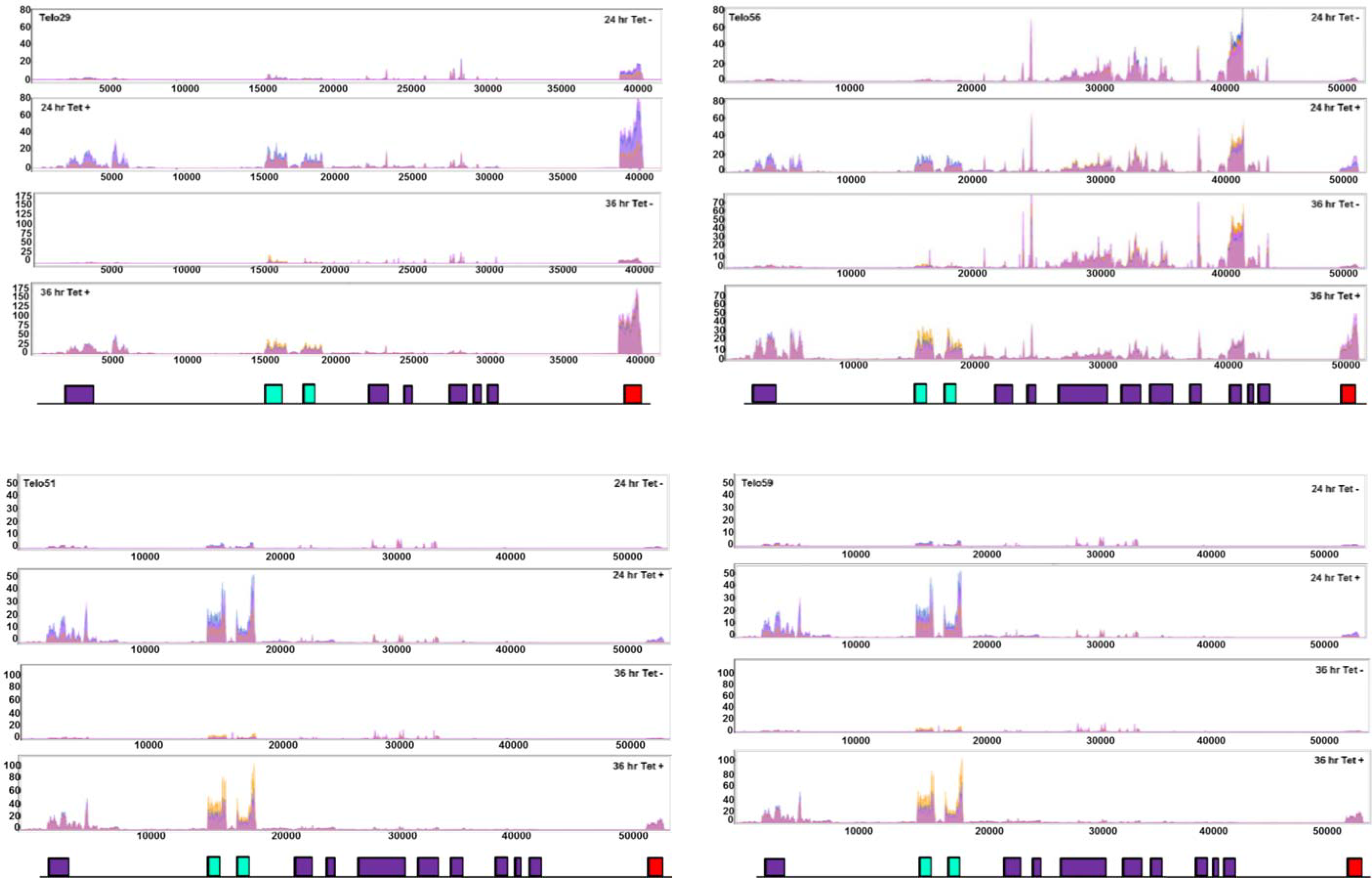
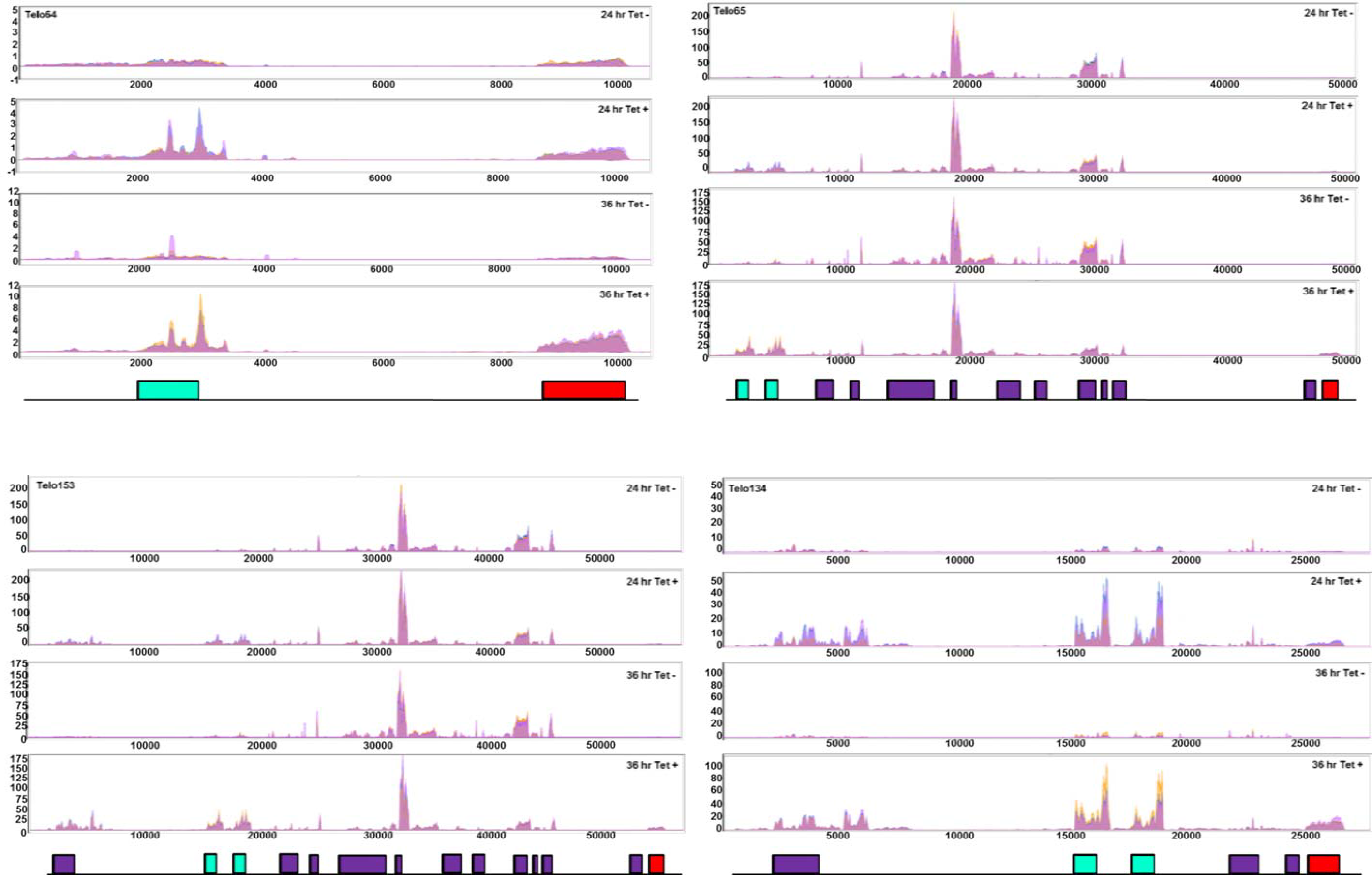
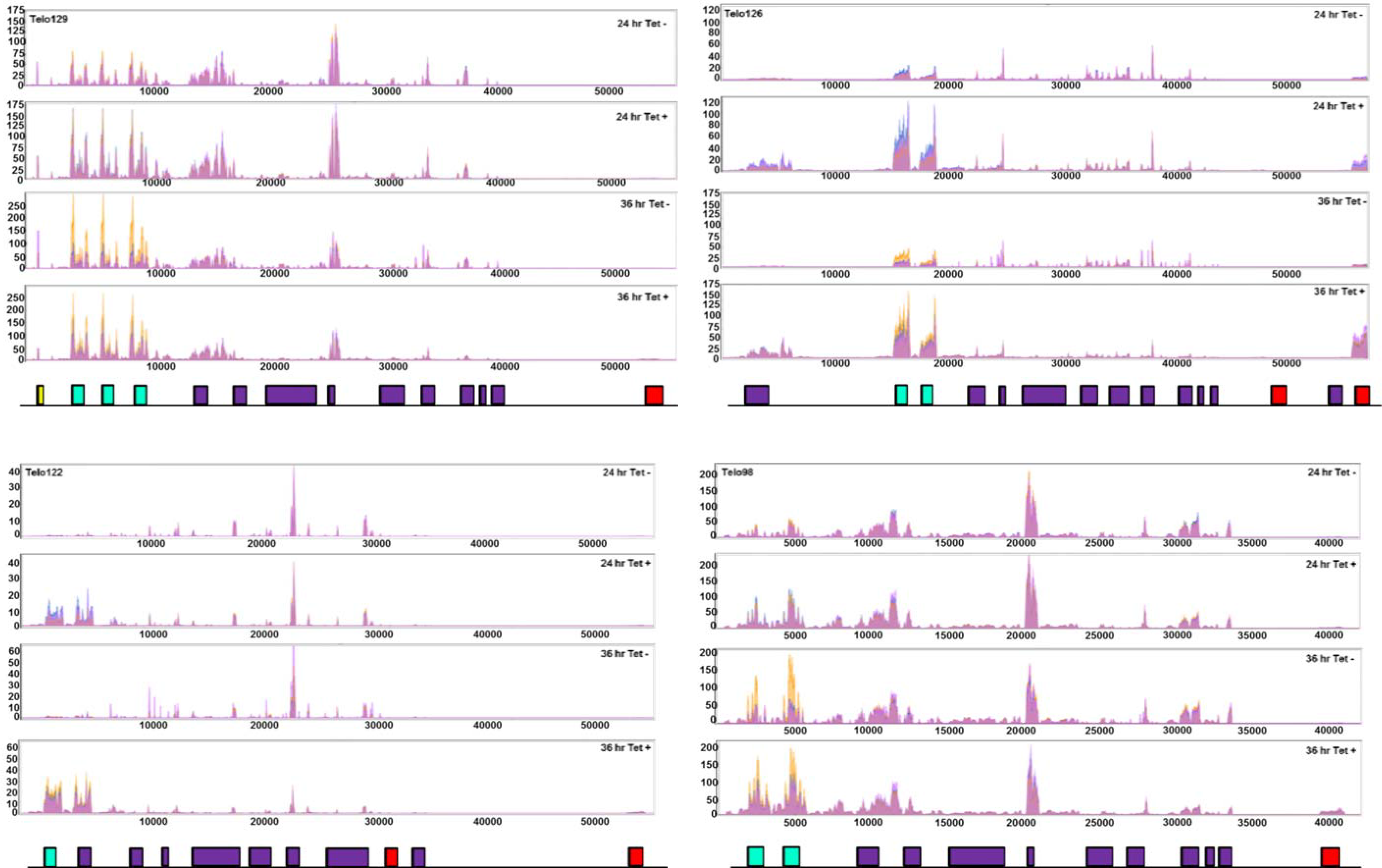
Loss of ATR impairs control of silent *T. brucei* bloodstream form VSG expression site transcription. Mapping of MapQ filtered RNAseq reads is shown across all silent VSG bloodstream VSG expression sites (except Telo4, which is shown in Fig.3 and Fig.S3) after 24 and 36 hrs growth with (T+) or without (T−) induction of TbATR RNAi; y axes show the number of reads that map relative to location in the transcription units, and data from three replicates are overlaid. ESAG6 and ESAG7 genes are shown in green, all other ESAGs are in purple, the VSG is in red.

**Fig.4 – supplementary figure 1.**
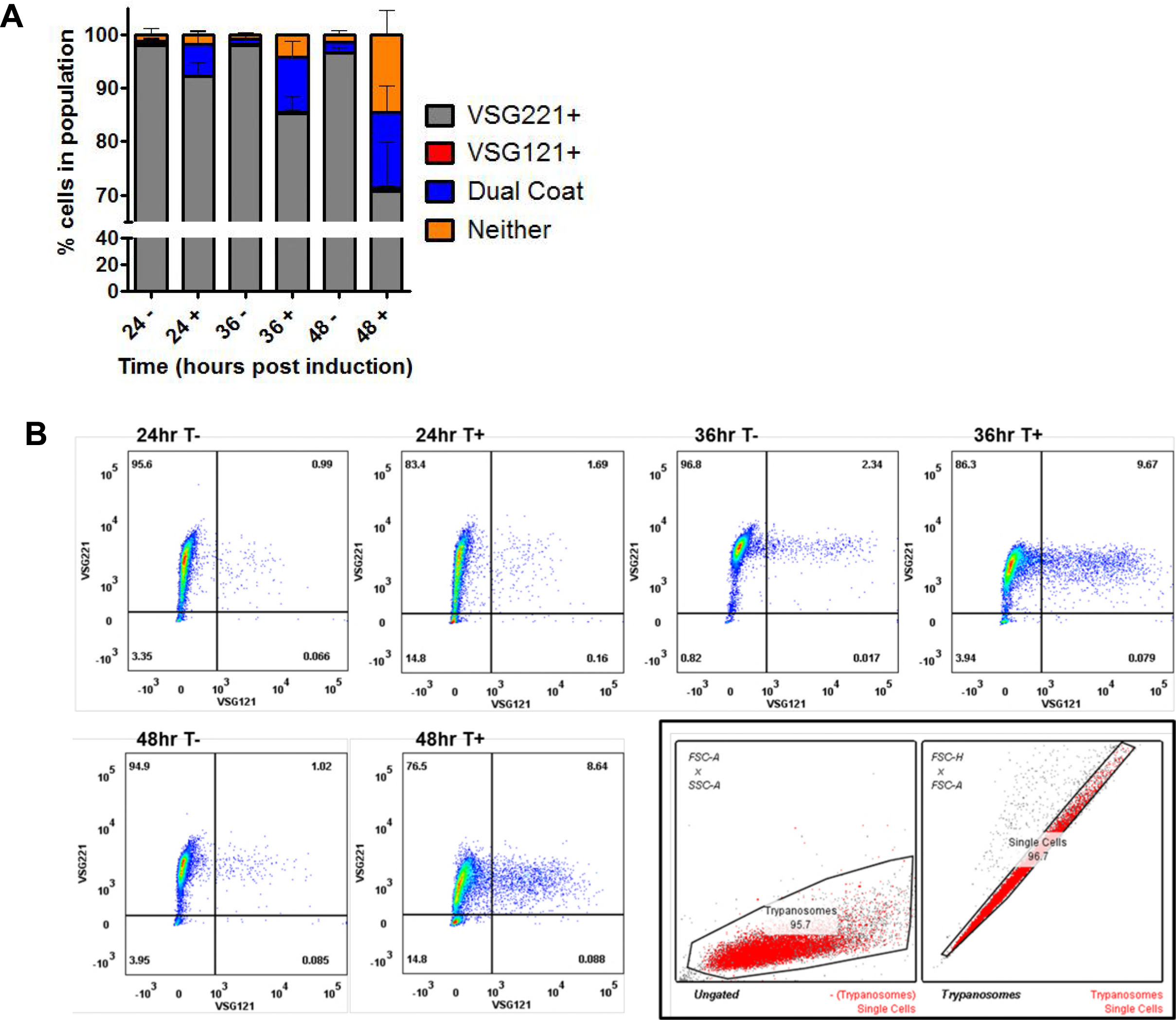
Loss of ATR in bloodstream form *T. brucei* results in changes in VSG coat expression. (A) Analysis of VSG expression by indirect immunofluorescence. Cells were collected at 24, 36 and 48 hours after TbATR RNAi induction (+) in CL2, or at the same time points without RNAi induction (−), and stained with anti-VSG221 and anti-VSG121 antiserum. Individual cells were scored for the presence of just one of the VSGs (VSG221+, grey; VSG121+, red), both VSGs (dual coat, blue), or neither VSG on their surface (orange); numbers are expressed as a percentage of the total population and error bars show ± SEM for three experimental repeats (>200 cells were counted at each time point in all experiments). (B) Analysis of VSG expression by flow cytometry. Cells in which TbATR RNAi had been induced (T+) or controls without induction (T−) were collected from CL2 after 24, 36 and 48 hours growth, stained with anti-VSG221 and anti-VSG121 antiserum and analysed by flow cytometry. Over 10,000 cells were analysed per sample and time point; representative date from one experiment is shown. The boxed plots detail the gating strategy used in the flow cytometry. To discriminate between healthy and dead cells, side scatter (SSC-A, linear) and forward scatter (FSC-A, linear) were plotted. Cells with high SSC-A signals or very low SSC-A signals suggest these cells are very granular (high) or debris (low). Wide gates were chosen to permit analysis of phenotypic changes following TbATR RNAi.

**Fig.5 – supplementary figure 1.**
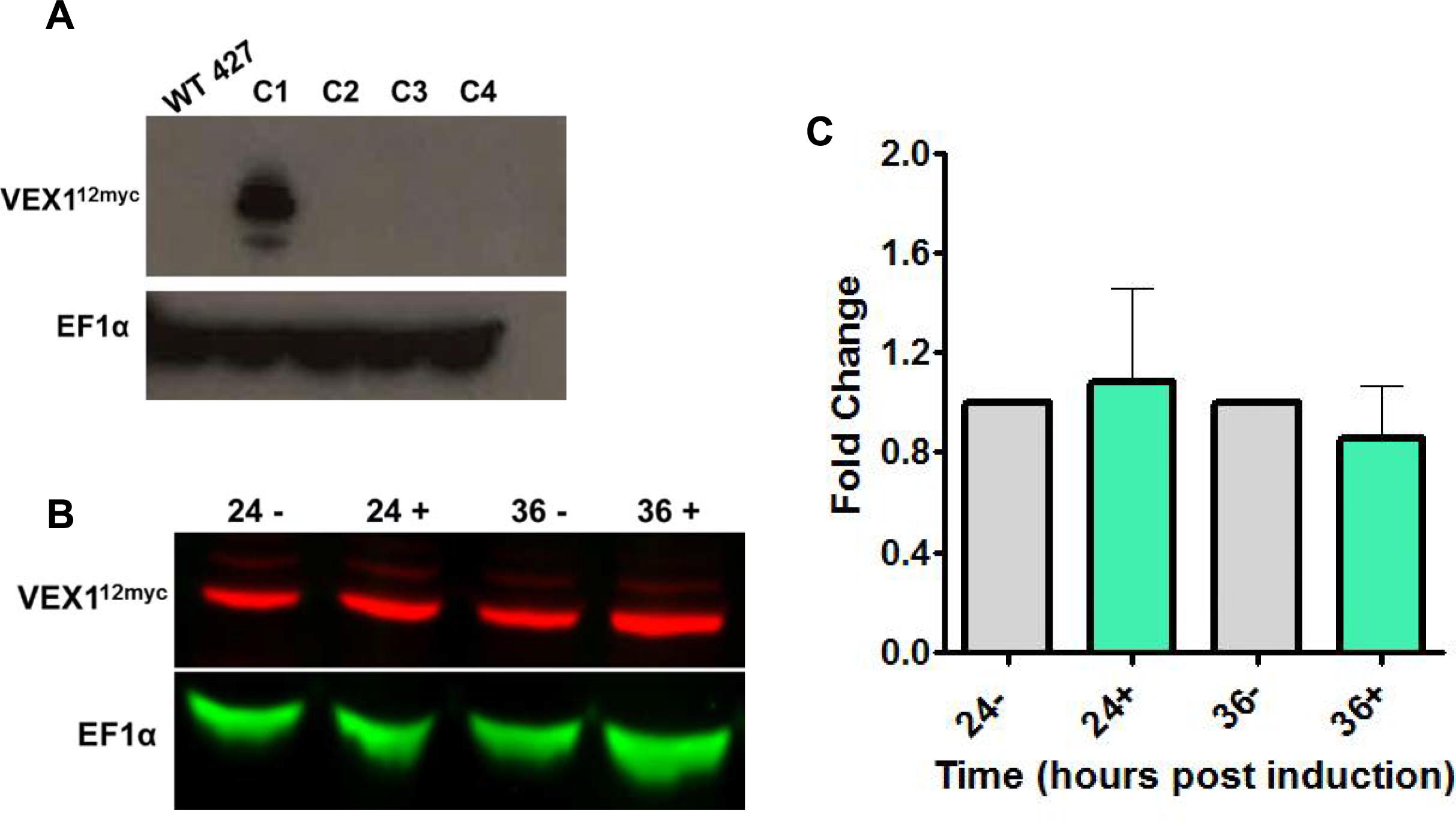
Loss of TbATR does not affect abundance of VEX1-12myc protein. (A) Transformant clones (C) putatively endogenously expressing VEX1 tagged with 12 copies of the myc epitope (VEX1-12myc) were generated, and expression of an appropriately sized protein that reacted with anti-myc antiserum was detected in one (C1); anti-EF1α antiserum was used as a loading control and blotting was performed with wildtype, untagged (WT427) cells. (B) Expression levels of VEX1-12myc (green) are shown by western blotting of whole cell extracts with anti-myc antiserum after 24 or 36 hours growth with (+) and without (−) addition of Tet; EF1α (red) serves as a loading control. (C) Fold-change in levels of VEX1-12myc protein after depletion by RNAi of TbATR at 24 and 36 hrs growth with (+) RNAi induction; values are shown relative to uninduced cells (set at 1.0) after normalisation using EF1α signal.

**Fig.5 – supplementary figure 2.**
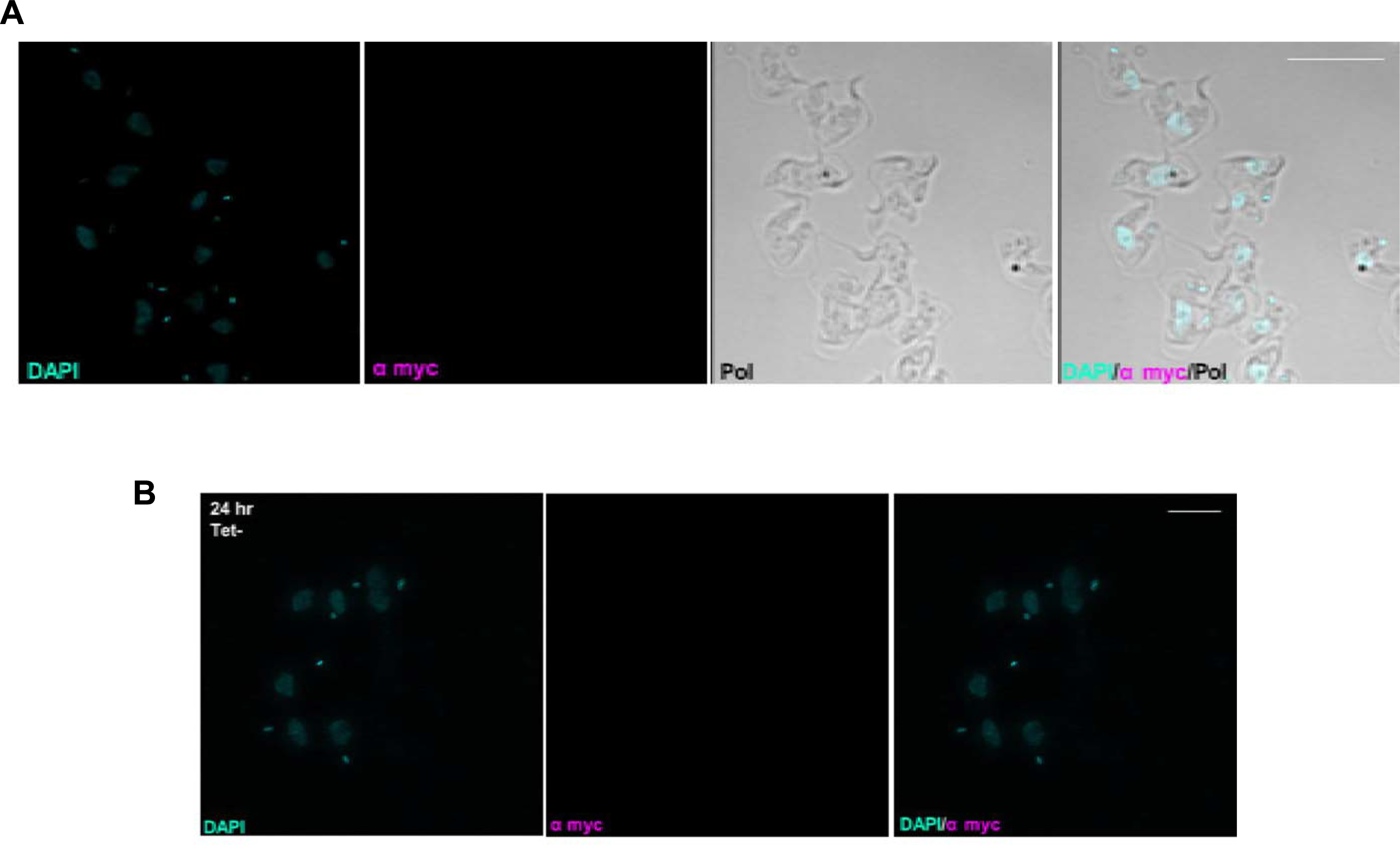
Untagged cells to control for non-specific anti-myc signal. (A) Untagged wild type (WT427) cells are shown stained with DAPI (cyan) and with anti-myc antiserum (magenta); the cell outline is shown by polarised light (Pol), and merged images are provided. Images were captured on a DeltaVision microscope; scale bar = 5 µm. (B) Untagged TbATR RNAi uninduced cells (teT−) are shown in the same way as in (A), but here images were captured on an Elyra super resolution microscope (no cell outline shown); scale bar = 5 µm.

**Fig.5 – supplementary figure 3.**
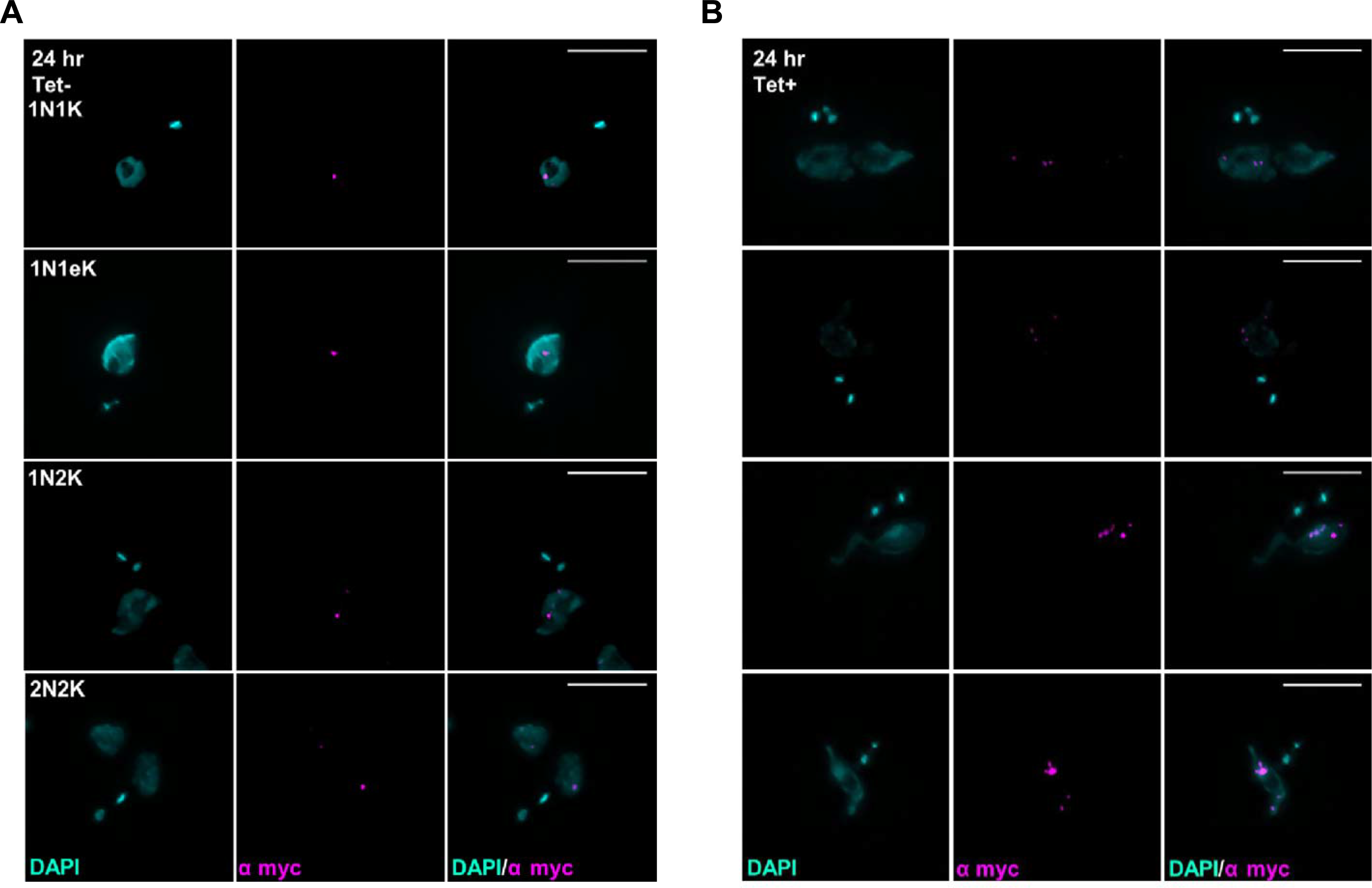
Localisation of VEX1-12myc. (A) Representative images of VEX1-12myc localisation (detected with anti-myc antiserum, magenta), in different cell cycle stages (determined by nuclear (N) and kinetoplast (K) configurations after DAPI staining, cyan), of uninduced (teT−) TbATR RNAi cells. (B) Representative images of VEX1-12myc localisation in TbATR depleted cells following 24 hrs RNAi induction (tet+). Images were captured on a DeltaVision microscope; scale bar = 5 µm.

**Fig.5 – supplementary figure 4.**
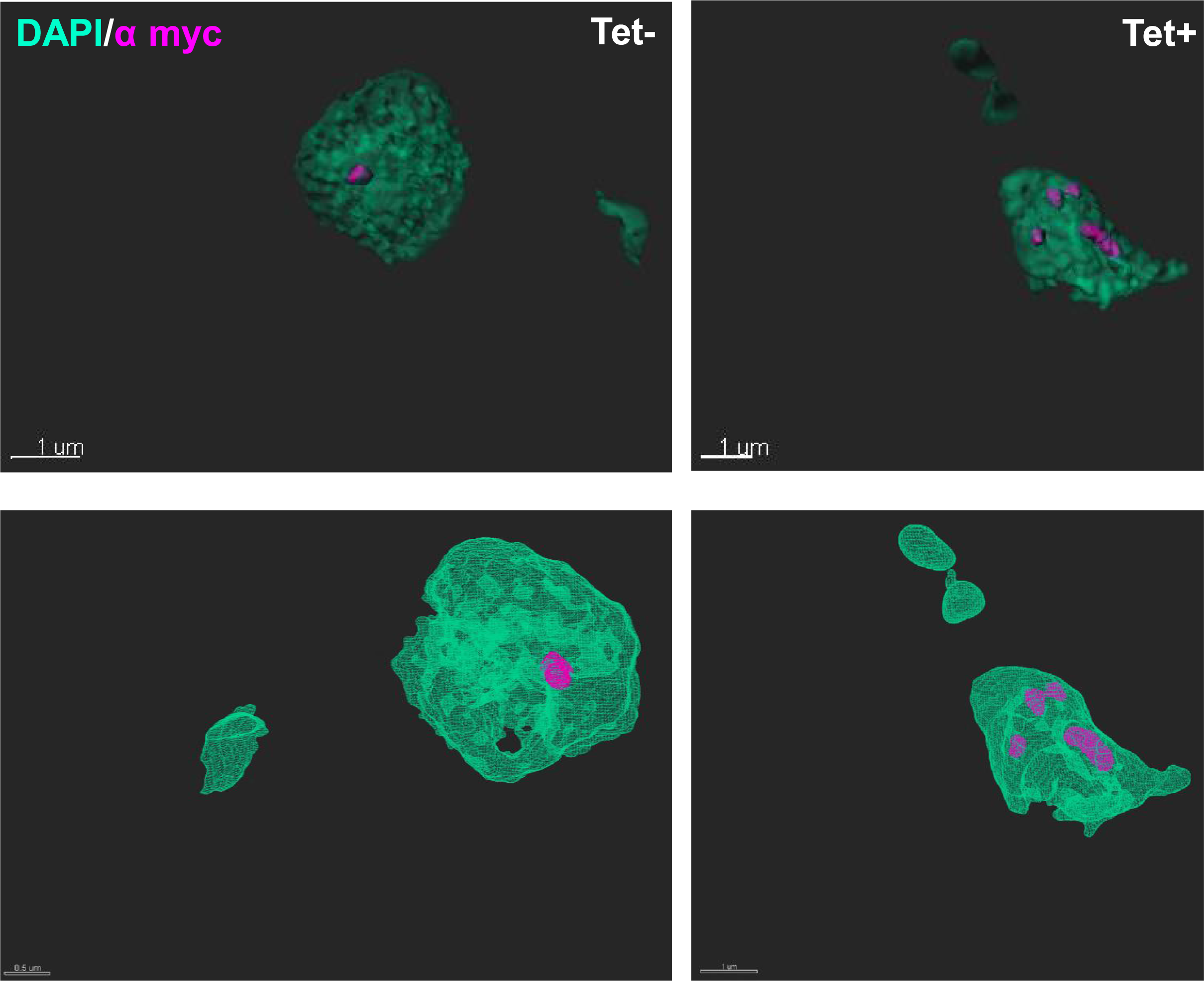
3D modelled images of VEX1-12myc localisation before (−) and after (+) RNAi depetion of TbATR. Z-stacked images were captured on an Elyra super resolution microscope and complied images generated using IMARIS software (V.8.2). Scale bars are as stated on the images. DNA were stained with DAPI (cyan), and VEX1-12myc detected with anti-myc antiserum (magenta).

**Fig.6 – supplementary figure 1.**
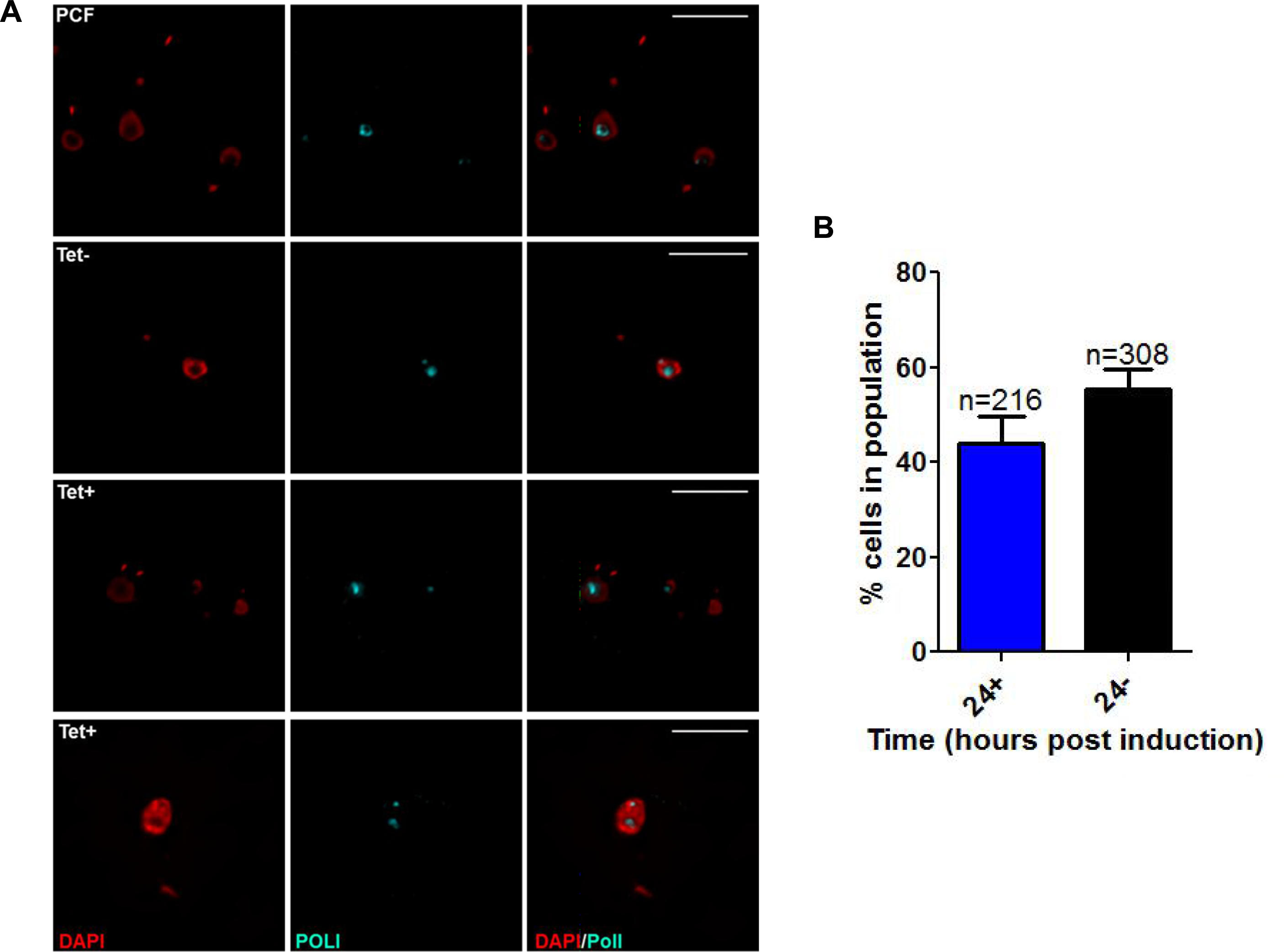
Loss of TbATR results in reduced numbers of ESB containing cells and aberrant RNA Pol I localisation. (A) Representative images of RNA Pol I localisation with (tet+) and without (teT−) induction of RNAi against TbATR in bloodstream form cells; procyclic form cells (PCFs) were used as a control. DNA was visualised by DAPI staining (red), and RNA Pol I detected using anti-Pol I antiserum (cyan). Images were captured on a DeltaVision microscope; scale bar = 5 µm. (B) Analysis of RNA Pol I and ESB presence after 24 hours growth with (tet+) and without (teT−) induction of RNAi against TbATR. Values represent the percentage of the total cells counted (n) in which nucleolar and distinct ESB staining could be discerned; error bars represent SEM from two independent experiments.

**Table S1.** RNAseq analysis of gene expression changes after 24 hours of TbATR RNAi.

**Table S2.** Go term enrichment analysis of gene cohorts found to be significantly up-regulated or down-regulated after 24 hours of TbATR RNAi.

**Table S3.** RNAseq analysis of bloodstream VSG expression site-localised transcripts found to be significantly up-regulated or down-regulated after 36 hours of TbATR RNAi.

